# Pythia: Non-random DNA repair allows predictable CRISPR/Cas9 integration and gene editing

**DOI:** 10.1101/2024.09.23.614424

**Authors:** Thomas Naert, Taiyo Yamamoto, Shuting Han, Melanie Horn, Phillip Bethge, Nikita Vladimirov, Fabian F. Voigt, Joana Figueiro-Silva, Ruxandra Bachmann-Gagescu, Fritjof Helmchen, Soeren S. Lienkamp

## Abstract

CRISPR-based genome engineering holds enormous promise for basic science and therapeutic applications. Integrating and editing DNA sequences is still challenging in many cellular contexts, largely due to insufficient control of the repair process. We find that repair at the genome-cargo interface is predictable by deep-learning models and adheres to sequence context specific rules. Based on *in silico* predictions, we devised a strategy of triplet base-pair repeat repair arms that correspond to microhomologies at double-strand breaks (trimologies), which facilitated integration of large cargo (>2 kb) and protected the targeted locus and transgene from excessive damage. Successful integrations occurred in >30 loci in human cells and in *in vivo* models. Germline transmissible transgene integration in *Xenopus*, and endogenous tagging of tubulin in adult mice brains demonstrated integration during early embryonic cleavage and in non-dividing differentiated cells. Further, optimal repair arms for single- or double nucleotide edits were predictable, and facilitated small edits *in vitro* and *in vivo* using oligonucleotide templates.

We provide a design-tool (Pythia, pythia-editing.org) to optimize custom integration, tagging or editing strategies. Pythia will facilitate genomic integration and editing for experimental and therapeutic purposes for a wider range of target cell types and applications.

---

**T**he precise and targeted integration of transgenes via CRISPR/Cas9 technology holds significant promise for applications in biotechnology and gene therapy^1^. However, it is paramount that genomic integrity is maintained to avoid unintended side-effects and that the integration technique is suitable for targeting the intended cell types^2,3^. Typically, CRISPR/Cas9-mediated integration relies on homology-directed repair (HDR), which necessitates large homology arms on either side of the cargo and is only active in proliferating cells, or on non-homologous end joining (NHEJ), microhomology-mediated end joining (MMEJ) or single-strand annealing^4^. However, NHEJ and MMEJ may result in unintended genomic alterations at transgene-genome borders, including genetic deletions within the surrounding genome or transgene, and the potential disruption of neighbouring genes^5,6^.

In humans, naturally occurring double-stranded breaks (DSBs) are typically repaired accurately, yet occasionally, inherently mutagenic MMEJ repair results in genetic errors. Microdeletion variants account for 20-25% of all clinically pathogenic sequence variants^7,8^. The majority of these mutations display a local sequence signature characteristic of deletions through microhomologies (µH) and are often three adjacent base pairs in length. Utilizing this natural MMEJ mechanism for frame retaining DSB repair of coding sequences offers tantalizing biotechnological opportunities. Predicting the repair process at the interface between the genome and transgene repair arms could thus lead to more rational design and more control over gene editing outcomes.

MMEJ as a repair mechanism for DSBs induced by CRISPR/Cas9 is conserved across a broad spectrum of organisms, ranging from Hydrozoa^10^ and plants^11^ to zebrafish^12,13^, *Xenopus*^14^ to humans^15,16^.

Such MMEJ repair occurs in a non-random fashion and is predictable by algorithms and deep learning models, such as InDelphi^17–19^. This predictability has been harnessed to establish programmable smaller^17^ and larger^20,21^ deletions after DSB repair, but never transgene insertions. While MMEJ-mediated approaches have been successfully employed for integration, for example GeneWeld^22^ and PITCh^23–25^, these did not offer control over gene editing outcomes at genome-transgene repair boundaries. On the other hand, prime editing’s effectiveness depends on the coordination of multiple components and is traditionally restricted to edits ranging from single to ∼50 base pairs (bp), rendering larger insertions inaccessible^26^. New tools that combine prime editors with serine integrases, such as TwinPE^27^, PASTE^28^ and PASSIGE^29^, have been shown to enable larger DNA insertions, yet leave a footprint, making them less suitable for protein tagging applications.

The original CRISPR/Cas9 system has been widely adopted in biotechnology and basic research. Here, we focus on the insertion of large cargo using this most frequently used genome editing technique and explore the predictable nature of double-strand break (DSB) repair mechanisms when introducing exogenous genetic material. Obtaining predictable outcomes in DNA editing could not only help predicting integration results using CRISPR/Cas9, but would also suggest that any transgenesis method relying on DSB repair could, to some extent, be programmable and controllable if the appropriate prediction model is applied. Here, we explored the potential for harnessing and mining deep learning models, pretrained on DNA repair outcomes, to develop the optimal rules for designing DNA repair arms, both for integration of large genetic cargo, as well as to establish small point mutations. This results in predictable editing outcomes driving intended edits and integrations, but not bystander mutations.

We used tandem repeats of exceptionally small homologies (<u>tri</u>-nucleotide repeating ho<u>mologies;</u> 3 bp tandem repeats, which we called trimologies), placed at the edges of transgene cassettes to facilitate on-target integration via MMEJ using CRISPR/Cas9. We find that DSB repair is non-random on the interface between the genome and trimology repair arms of large transgenic cassettes (>2.5kb), *in vitro* and *in vivo*, unveiling an unanticipated predictability. Moreover, trimology repair arms safeguard the boundaries during integration, precluding extensive DNA trimming. We further deducted optimal design rules and show trimology integration to be effective in cell contexts where HDR is largely ineffective, such as rapidly cycling vertebrate embryos (*Xenopus*) and adult post-mitotic mouse neuronal cells. Finally, we extend this novel notion of predictability to rational design of small repair templates for desired point mutations at permissive loci with single stranded DNA (ssODN) donor templates.

## Cas9 integration with donor templates is non-random and predictable

Endogenous DNA repair outcomes following CRISPR/Cas9 induced double-stranded breaks (DSB) are non-random and can be predicted based on the local sequence context^15–18^. We explored, if one of such prediction algorithms, InDelphi^17^, could also predict editing outcomes at the interface between endogenous DSB edges and exogenous donor DNA. When the InDelphi model predicted a micro homology (µH) mediated 4 bp deletion as the major editing outcome of an example sequence (Fig. 1a), adding a 3 bp sequence initially present on the left side of the cut to the sequence right of the cut, pivoted the most frequent predicted outcome towards a 3 bp deletion. This effectively removed the inserted 3 bp µH, overruling the previously dominant 4 bp deletion in this particular example. Further repeating the 3 bp sequences in tandem (dubbed trimology repeats) increased the amount of predicted editing outcomes that use an inserted artificial µH from 52% to 62% (Fig. 1a). Extending the *in silico* simulation of the InDelphi models to 250.000 putative gRNAs target loci on human chromosome 1 revealed an increase in artificial µH usage for DNA repair with increasing number of tandem repeats, which plateaued at five tandem repeats (Fig. 1b, S1). Further increases in tandem repeats beyond five showed only minor improvement in µH usage. However, the local sequence context strongly influenced the use of µH tandem repeats for each individual gRNA (Fig. 1c), suggesting that the optimal design needs to be computed for each gRNA and its surrounding genomic sequence, when predicting µH-mediated repair outcomes.

**Fig. 1.**
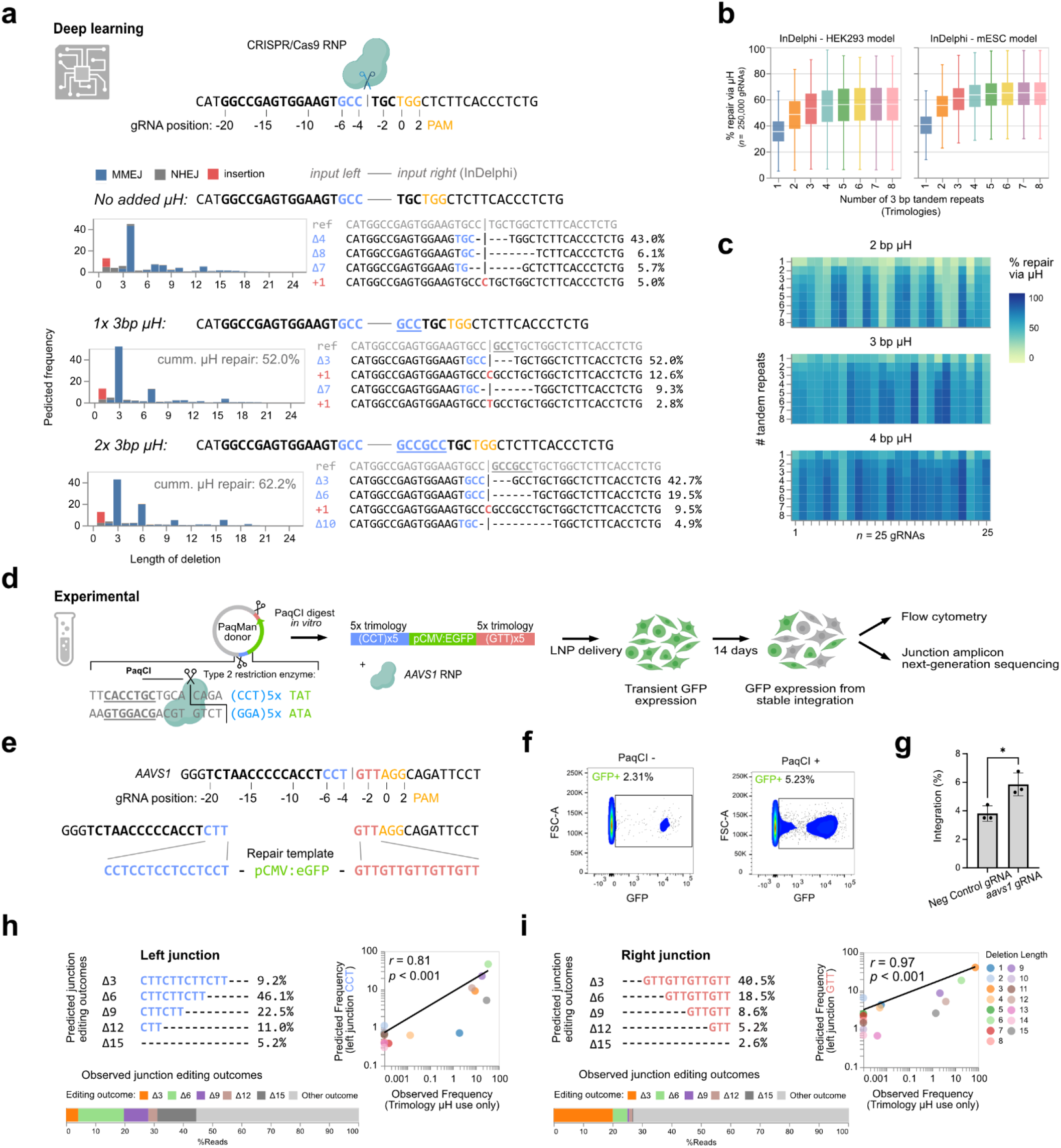
Modelling predicted gene editing outcomes using InDelphi while providing synthetic microhomologies. **(a)** Deploying InDelphi (HEK293T) on a synthetic DNA, the predicted editing outcomes are shown. Adding tandem repeats of the bases left of the CRISPR/Cas9 cut site, to the right of the cut affected the predicted editing outcomes. Cumulative µH repair is defined as the percentage of editing outcomes that mobilize (delete) a synthetic µH during repair. **(b)** Modelling of expected editing outcomes across 250,000 distinct gRNAs target sites across human Chr1, when adding the three base pairs (trimologies) flanking the left site of the CRISPR/Cas9 cut site either as a single repeat (1x) or as tandem repeats (2-8x). The % of repair via µH usage is shown. **(c)** Heatmap highlighting the expected % of repair via µH as a function of the length of µH and the number of tandem repeats for 25 gRNAs, demonstrating that there is a sequence context-specific optimal solution for maximizing the percentage of µH repair outcomes. **(d)** Schematic of the experimental setup: PaqCI digestion releases the linear dsDNA donor, which contains 5X trimology arms, and is co-delivered with RNP targeting *AAVS1*. **(e)** Sequence of the target locus and trimology containing repair arms. **(f)** After 14 days, Flow cytomety after 14 days indicates an increase in stable integration in cells transfected with the linear dsDNA template. **(g)** Quantification of integration efficiency of *AAVS1* gRNA in comparison to a negative control gRNA (t-test, p<0.05, n=3). **(h-i)** The Indelphi-HEK293T model accurately predicts the observed frequency of distinct editing outcomes in the trimology arms at both junctions.

Next, we experimentally investigated whether InDelphi predictions of junctional products at the interface between endogenous DSB edges and exogenous donor DNA would facilitate CRISPR/Cas9-mediated knock-in. For this, the *AAVS1* landing site was targeted in HEK293T cells (Fig. 1d). We added five tandem repeats of 3 bp microhomology arms (5x trimologies) to the left and right of the donor cassette, thus matching the sequence context left and right of the *AAVS1* cut site (Fig. 1e). Trimology repair arms were chosen to investigate the scarring patterns that occur during DNA repair and validate the predictability of DNA repair at genome-transgene borders while enhancing the expected frequencies of frameshift retaining editing outcomes (3 bp repeats). To easier customize the donor edges without undesired 5’->3’ overhangs, we added two PaqCI type IIS endonuclease restriction sites invertedly flanking the donor cassette (pCMV:eGFP) and the trimologies for *in vitro* release of linear donor DNA from plasmids (PaqMan plasmids, Fig. S2a). PaqMan linearization facilitated on-target genomic integration (5.2% GFP+), while nonlinearized PaqMan plasmid donor merely resulted in random integration (2.3% GFP+) (Fig. 1f), demonstrated by boundary PCR analysis (Fig. S2b). On-target integration further only occurred with an *aavs1* targeting RNP and never with control RNP (gRNA target site not present in the human genome) (Fig. 1g, Fig. S2c).

Using trimology repair arms provided us with a unique way to sample the distribution of editing outcomes at the interface between endogenous DNA and exogenous cargo. By observing which microhomologies were used at which percentage, it was possible to determine whether junctional editing outcomes were correctly predicted by the InDelphi model. Targeted amplicon sequencing of the boundary PCR products revealed that the rate of trimology usage after DNA integration observed experimentally correlated well with the InDelphi predictions at the left (*r*=0.81, *p*<0.001) and right junction (*r*=0.97, *p*<0.001) (Fig. 1h-i, Fig. S3a), validating our *in silico* models. Further this revealed that on the left junction boundary, 73% of the reads did not trim into the genome. Of these 63% (46% of total reads) also did not trim into the transgene (Fig. S3b-c). Similarly, 78% of the reads on the right junction did not trim into the genome, 55% of these (43% of total) also did not trim into the transgene. Interestingly, the most common genetic lesion after trimming-free integration was the loss of one or more of the trimologies in the repair arms (45% total reads left, 28% right) (Fig. S3b-c).

Thus, merely 3 bp of µH were sufficient to mobilize DNA donor arms during CRISPR/Cas9 knock-in. Taken together, we show that Cas9-mediated MMEJ integration is non-random and predictable.

## Trimology arms safeguard the genome and integration efficiency is influenced by local sequence context

Next, we benchmarked our methodology to NHEJ-mediated gene cassette knock-in, such as the HiTi methodology which does not utilize homology arms^30^. We generated PaqMan plasmid donors containing either zero or four trimologies matching the *AAVS1* target site and investigated integration efficiencies and junctional gene editing outcomes (Fig. 2a, S2d). There were no detectable differences in integration efficiencies when using either zero (NHEJ - 9.3%) or four (10.7%) trimologies in HEK293T cells (Fig. 2b) (p > 0.05). When NHEJ was used, however, amplicon sequencing revealed extensive deletions of the genome (95% of reads). All remaining reads showed substantial trimming of the transgene cassette (Fig. 2c-d). In contrast, adding trimology repair arms decreased DNA trimming both into the genome and on the repair cassette with over 50% of reads free from any deletions in either direction. This demonstrated that adding trimology arms to the donor templates had a protective effect on extensive DNA end-trimming.

**Fig. 2.**
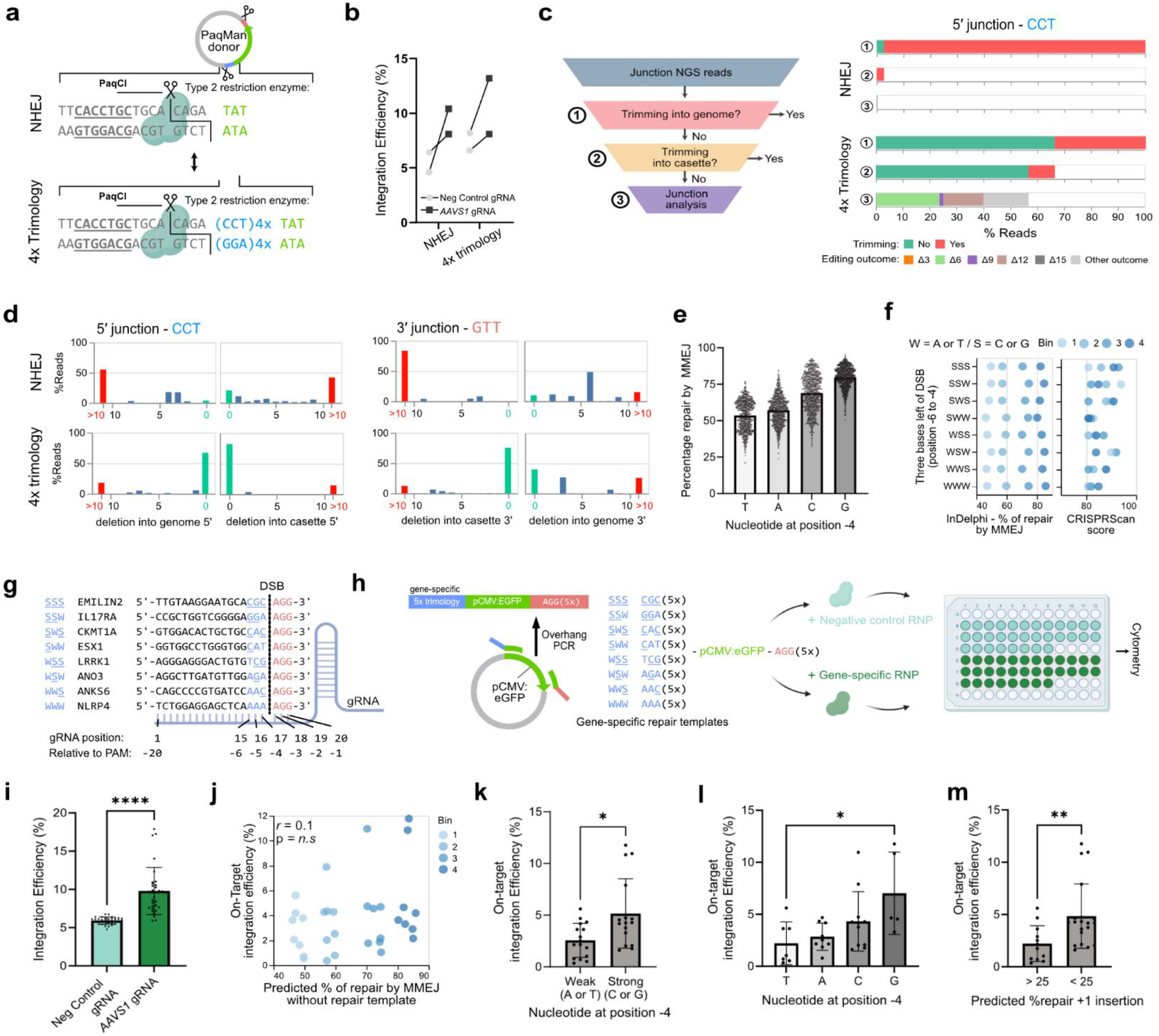
Trimology arms protect the genome and the relationship between integration efficiencies and local sequence context. **(a)** Schematic representation of the experimental setup comparing NHEJ integration (no repair arms) to 4 tandem repeat trimology integration using PaqMan plasmids. **(b)** Comparision of trimology integration and NHEJ integration efficiencies. (t-test, p>0.05, n=2)) **(c)** Visualisation of genome editing outcomes on both genome-transgene junctions showing % of reads Fig. 2 (Cont.) |that trimmed the genome (1), reads that trimmed the cassette (2), and specific editing outcomes of reads that trimmed neither the genome nor the cassette (3). **(d)** Quantification of genome editing outcomes on both genome-transgene junctions demonstrating that in NHEJ leads to extensive trimming, while 4X trimology arms protect both the genome and transgene cassette. **(e)** In the absence of exogenous DNA, *in silico* modelling predicts the the nucleotide at position -4 will influences the percentage of repair outcomes that is expected to be driven by MMEJ. (total *n*=10,813,171 - plotted random subselection of 500,000 datapoints). **(f-g)** 32 gRNAs designed to target coding exons of non-essential genes with four in each of eight classes covering all possible permutations of strong (G or C) and weak (A or T) bases in three bases left of the DSB. Each class was composed of 4 gRNAs binned across InDelphi-predicted % of repair by MMEJ and had similar expected on target efficiencies (CRISPRScan scores). **(h)** For each gRNA, a distinct dsDNA repair template was generated with 5x trimologies matching either the gRNA-specific context left of the DSB, and 5x trimologies matching the “AAG” right of the DSB. These were delivered either with non-targeting control RNP (top) or gene-targeting RNP (bottom) to HEK293T cells. **(i)** Integration efficiencies at day 14, determined by flow cytometric quantification of GFP+ cells. (Mann-Whitney, p<0.0001, n=32) **(j)** Scatter plot of InDelphi predicted % of repair by MMEJ at the endogenous locus (in the absence of exogenous DNA) and the on-target integration efficiency (normalized for random background integration). **(k)** Quantification of on-target integration efficiencies comparing the presence of a strong or weak base at position -4, just left of the DSB (Mann-Whitney, p<0.05, n=16). **(l)** On target integration efficiencies depending on each base at position -6 is highest with guanine (Kruskal-Wallis, p<0.05, n=8). **(m)** InDelphi modelling of the junction product between the sequence left of the DSB and the dsDNA donor. A higher percentage of predicted editing outcomes that have a +1 insertion will result in a lower on-target integration efficiency (Mann-Whitney, p<0.01). All error bars are S.D.

Next, we tested if the nucleotide composition of trimology arms affected their integration efficiency. Initially, we performed an *in silico* simulation with the InDelphi-HEK293 model, predicting editing outcomes for >10 million gRNAs across the entire human genome. Notably, the model forecasted variations in repair outcomes driven by microhomology composition, particularly linked to the nucleotide at position -4 (counting the NGG PAM as nucleotides 0-2) (Fig. 2e). A G at position -4 was predicted to enhance integration over a C, A or T, which was least favourable. This effect remained consistent across simulations using either NGG or NAA PAM sequences (Fig. S4-5), suggesting that it was independent of PAM composition and constraints. Notably, no similar effects were noted for any other position in the gRNA (Fig. S4-5). Our *in silico* findings indicated that the nucleotide at position -4, thus located immediately to the left of the CRISPR/Cas9 induced DSB, could influence the proportion of DSB repair that utilizes microhomology-mediated mechanisms and could be a parameter to improve trimology integration.

To experimentally test this hypothesis, we employed CRISPR/Cas9 to target 32 genes in HEK293T cells and co-delivered target-specific repair templates that included five trimologies. To avoid a potential negative selection effect, we chose genes previously identified as non-essential^31^. We ensured that the gRNAs had similar predicted on-target efficiency and a balanced distribution across different GC-contexts (Fig. 2f). To directly assess whether the nucleotides at position -7 to -4 influence trimology integration, we only considered gRNA target sites with “AGG” at nucleotides -3 to -1. The 32 targets were chosen to fall into one of eight distinct classes, each class representing a distinct combination of strong (G or C) or weak (A or T) bases at positions -4 to -7 (*n*=4 per class) (Fig. 2f-g). Within each class, we selectively identified gRNAs and binned them according to predicted MMEJ repair usage. Of note, target-specific repair templates, incorporating five trimologies, were generating by straightforward overhang PCR (Fig. 2h).

Across all 32 targets, we observed a median 1.6-fold increase in integration efficiencies comparing on-target RNP to negative control RNP (median on-target integration of 3.61%, p<0.0001, *n*=32) (Fig. 2i). There was no correlation between on-target integration efficiency and InDelphi predicted MMEJ repair at the DSB (no exogenous repair templates in modelling) (Fig. 2j). This indicated that the presence or absence of microhomologies surrounding the DSB did not directly influence integration efficiencies of exogenously provided repair cassettes. Interestingly, gRNAs that had a strong base (G or C) at nucleotide -4 drove integration at a median 1.8-fold more efficiently than those with a weak base at that position (*p*<0.05) (Fig. 2k). We found a hierarchical trend at nucleotide -4, where G (7% ±4%), C (4.3% ±2.9%), A (2.8% ±1.3%), and T (2.2% ±2.1%) influenced the use of microhomology mediated integration rates. (Fig. 2l). Notably, this distribution completely matched the predicted distribution (compare to Fig. 2e). Next, we used InDelphi to predict gene editing outcomes at the left junction between the endogenous locus and the cargo template. We observed a moderate inverse correlation between integration efficiencies and the percentage of repair predicted to be a +1 insertion (*r*=-0.512, *p*<0.01) and between integration efficiencies and the predicted percentage of perfect repair products (defined as having used one trimology) (*r*=0.51, *p*<0.01) (Fig. S6). Finally, we observed a median 2.2-fold higher rate of integration efficiency at junction events where InDelphi predicted the editing outcomes to be <25% +1 insertions than >25% +1 insertions (*p*<0.01) (Fig. 2m). Based on these observations we propose the following for selecting gRNAs for optimal trimology integration: (1) a G nucleotide at position -4, (2) low rate (<25%) of predicted editing outcomes with a +1 insertion and (3) a high percentage of predicted editing outcomes that utilize trimology arm µHs. Collectively, our experimental findings confirm that deep learning based predictions can enhance trimology integration outcomes and establish rationality in designing optimal integration strategies.

## Trimology integration of large cargo *in vivo* at the *h11* landing site of *Xenopus tropicalis*

Existing transgenesis methods (I-SceI^32^ and REMI^33^) to generate reporter lines in *Xenopus* are limited to random and multiple integration events. We identified a conserved *h11* locus^34^ on chromosome 1 of the *X. tropicalis* genome, in the intergenic sequence between *drg1* and *eif4enif1*, as a potential locus for stable transgene integration. *X. tropicalis h11* flanking gene models showed direct synteny with chicken, pig, human and rat orthologs (Fig. S7a)^35^. We identified two gRNAs (*h11*-α and *h11*-β), spaced 767 bp apart, and verified efficient editing activity (*h11*-α: 91% ± 9%, *h11-*β: 80.3% ± 2%) (Fig. S7b-f). Linear donor DNA containing four trimology repair arms corresponding with *h11*-α on the left and with *h11*-β on the right was generated by overhang PCR of a plasmid encoding CMV:eGFP (Fig. 3a). We co-injected *h11*-α and *h11*-β Cas9 RNP together with the trimology donor template into both blastomeres of two-cell stage embryos (Fig. 3a). We consistently observed eGFP expression across different developmental stages, indicative of stable integration events (Fig. 3b, S8d). PCR analysis of embryo pools (n=25) revealed CRISPR/Cas9-mediated deletion of DNA between the *h11*-α and *h11*-β target sites (Fig. S8a-b) and PCR junction fragments indicative of exogenous cassette integration in *h11* (Fig. 3c, Fig. S8c, e). To further simplify, using only one gRNA *h11-* α, one-cell stage just-fertilized embryos were targeted with a CMV:eGFP transgene containing four trimology repeats (Fig. 3d). 3% of embryos (*n*=4/134) were half-transgenics, suggesting that integration occurred at the two-cell stage (Fig. 3e), which could be clearly distinguished from embryos with a mosaic expression pattern (Fig. 3f). Junction products indicative of on-target integration could be amplified for half-transgenics, but never for embryos with mosaic eGFP expression (Fig. 3f). Sequencing confirmed the usage of trimologies for MMEJ-mediated repair (60%; *n*=5) (Fig 3g).

**Fig. 3.**
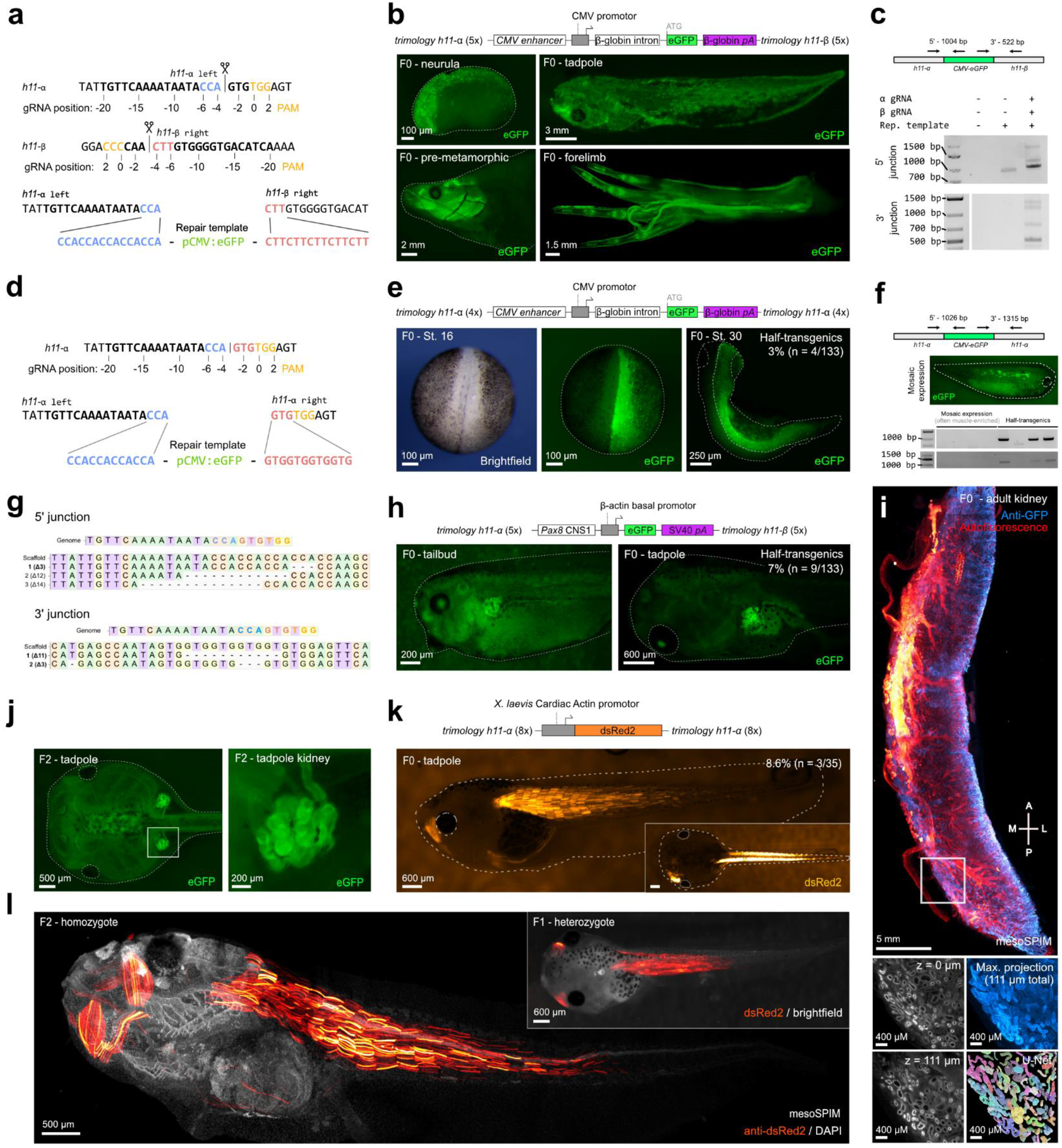
Trimology integration at stable landing site *h11* in *X. tropicalis* with germline transmission. **(a)** Schematic of the CRISPR/Cas9 integration strategy. **(b)** Mosaic, but stable GFP expression after trimology integration of a pCMV:eGFP in F0 founders at various developmental stages. **(c)** Detection of PCR products demonstrating on-target integration into the *h11* locus. **(d)** Schematic of the CRISPR/Cas9 integration strategy, using only the *h11-α* RNP. **(e)** Unilateral non-mosaic GFP expression in F0 founders due to pCMV:eGFP integration into the *h11* locus at the two cell-stage (half-transgenics). **(f)** Non integrative mosaic expression pattern in muscle cells. Junction PCR analysis shows that this represent merely transient expression as correct junction products can only be detected in half-transgenic animals shown in e. **(g)** Sequencing of junction products reveals usage of trimology repeats in 60% of reads (n=5). **(h)** Tissue-restricted expression pattern of *pax8*-CNS1:eGFP knocked-in at the *h11-α* and *h11-β* locus in the F0 generation via 5 trimologies is observed in 7% of the injected embryos (*n*=133). **(i)** Benchtop mesoSPIM whole-organ imaging of a kidney from an adult F0 Pax8-CNS1:eGFP founder, confirming stable integration and expression in renal tubules amenable for U-Net based segmentation. **(j)** Reporter expression in the embryonic kidneys of the F2 generation. **(k)** Tissue-restricted expression pattern of CarAct-dsRed knocked-in at the *h11-α* locus in the F0 generation via 8 trimologies is observed in 8.6% of injected embryos (*n*=35). **(l)** Benchtop mesoSPIM imaging of a F1 and F2 CarAct-dsRed knock-in animals revealing stable and strong tissue-restricted transgene expression.

One application of trimology integration is characterizing cis-regulatory sequences by integrating a candidate non-coding element with a minimal promoter and analysis of reporter expression levels and tissue-specificity^36^. However, this is only feasible when the number and site of integration can be controlled^37,38^. Therefore, a pax8-CNS1:eGFP construct was targeted to the *h11* locus using trimology integration^39^. In 7% of the injected embryos (n = 9/133), eGFP expression was observed in the pronephros, otic vesicle, and, to a lesser extent, the neural crest, consistent with the described activity of the cis-regulatory element (Fig. 3h, Fig. S9a). Trimology integration resulted in stable and persistent transgene reporter activity observed in the kidney tubules of adult F0 frogs (Fig. 3i, Movie S1). Germline transmission was confirmed in 50% (*n=6*) of F0 founder animals crossed with wild-type and 33%±12% of F1 embryos exhibited tissue specific GFP expression (Fig. 3j, Fig. S9b-c, Movie S2).

The trimology integration approach was further validated *in vivo* by integrating a *Xenopus* cardiac actin-dsRed2 (CarAct:dsRed2) reporter cassette (Fig. S9d)^37,40^. Strong non-mosaic muscle-specific dsRed2 expression was observed in 8.6% (*n* = 3/35) of the F0 animals (Fig. 3k). Germline transmission was successfully confirmed in both assessed founder animals (one male and one female), with each showing transmission rates of 10.5% and 45.5%, respectively, in dsRed2+ F1 embryos (Fig. S9e, Movie S3). In F2 homozygotes, we confirmed tissue specific dsRed2 activity in myotomes (Fig. 3l, Movie S4) and found single-copy integration of the reporter construct at *h11* (Fig. S9f). Taken together, we successfully achieved single-copy integration at the *h11* landing site in *X. tropicalis* of multiple donor templates without position effects or generational silencing.

## Trimology-mediated *in vivo* fluorescent tagging of Tubb2a in mice

Traditional HDR is ineffective in non-proliferating cells, but NHEJ-dependent HiTi is frequently utilized^30^. Thus, we asked whether addition of trimology repair arms could concurrently activate NHEJ and MMEJ, potentially increasing the proportion of in-frame tagged repair products facilitated by the frame retentive trimology design.

For this purpose, we targeted the 3’ end of *Tubulin Beta 2A Class IIa* (*Tubb2a*), a neuronal-specific tubulin localising to both axons and soma^41^, for in-frame eGFP tagging. We performed *in vivo* transduction by adeno-associated virus (AAV) into adult mouse brains (Fig. 4a-b). One AAV carries the Cas9, while the other carries *Tubb2a* targeting gRNA, promotorless eGFP for in-frame *Tubb2a* tagging and an ubiquitous promotor driving mCherry for assessing cargo delivery. Three weeks post-transduction, eGFP positive neuronal cells, featuring eGFP-tagged Tubb2a protein driven from the endogenous *Tubb2a* promotor, were observed via classical histology (Fig. 4c) and in volumetric mesoSPIM imaging of a whole-mount mouse brain optically cleared by modified wildDISCO^42^ (Fig. 4d, Movie S5). Cortical and hippocampal neurons with in-frame Tubb2a tagging were seen exclusively in virus-infected areas and displayed eGFP expression along their projections.

**Fig. 4.**
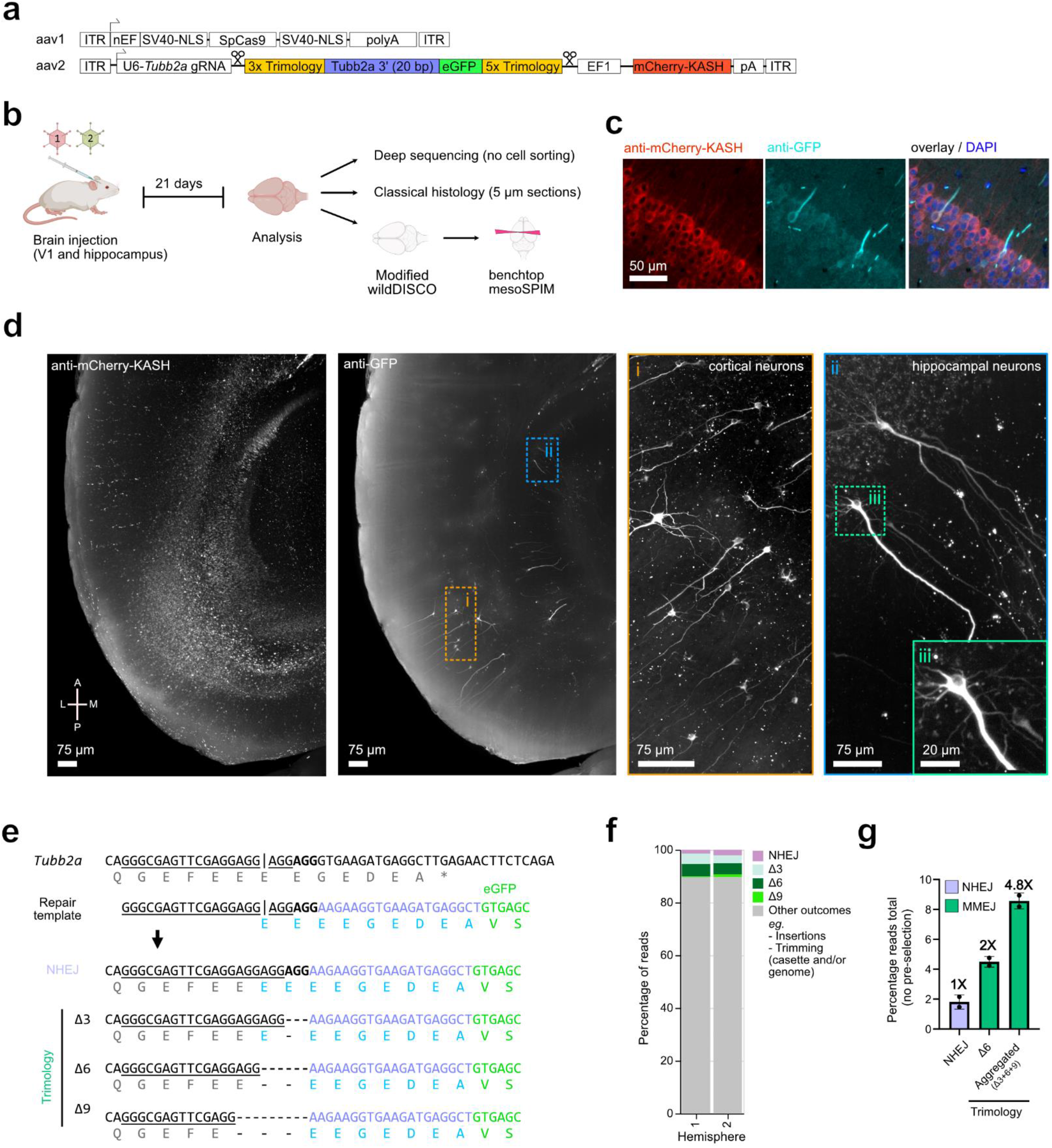
Endogenous fluorescence tagging of tubb2a *in vivo* in adult mouse brains via trimology integration. **(a)** Schematic of adeno-associated viral constructs for targeted eGFP knock-in at the 3’ CDS of *Tubb2a*. **(b)** Schematic of the experimental setup and subsequent analysis. **(c)** Histology of brain tissue and immunofluorescence detects eGFP-tagged Tubb2a in individual neurons. **(d)** Benchtop mesoSPIM light-sheet imaging of wildDisco-cleared whole mouse brain shows eGFP tagged Tubb2a in cortical and hippocampal neurons. **(e)** Sequence of the targeted *Tubb2a* locus (gRNA underlined, PAM in bold), the repair template and possible NHEJ and Trimology editing outcomes. **(f)** Summary of integration outcomes using NGS reads spanning *Tubb2a*-eGFP amplified from two mouse hemispheres. **(g)** Frequency of in-frame reads of *Tubb2a*-eGFP detecting either NHEJ or trimology integration outcomes as defined in panel e.

Next, we deep-sequenced the expected *Tubb2a*-eGFP junction site in two independently injected mouse hemispheres. Compared to earlier studies^30^, we did not preselect for cells expressing eGFP, thus getting an unbiased view of the gene editing outcomes at the expected junction site. HiTi repair outcomes, defined as a scarless NHEJ-mediated fusion between genome and transgene, only occurred in 1.8+-0.5% percent of the reads (Fig. 4e-f). While we indeed detected NHEJ-mediated gene tagging, it accounted for only a small subset of editing outcomes. Importantly, MMEJ-mediated mechanisms were active in post-mitotic cells as we observed 8.6+-0.5% gene editing outcomes that had utilized trimology dependent repair. As predicted by InDelphi, the most common editing outcome is a deletion of 6 nucleotides occurring at a frequency of 4.5+-0.4%. As such, our design strategy and use of trimology repair arms increased the number of reads containing in-frame mutations by 4.8-fold and of scar-free gene tagging by 2-fold, when compared to reads containing HiTi/NHEJ outcomes (Fig. 4g).

Taken together, trimology-mediated integration dramatically increased the efficiencies of in-frame gene tagging in mouse brains by engaging not only the NHEJ but also the MMEJ repair pathways.

## Pythia editing: precise genome rewriting via rational design of ssODN repair templates driving predictable DNA repair

Because junctional products of trimology integration were successfully predicted, we asked if the predictive power of InDelphi could also be used to design more customized editing strategies. For example, would the model be able to predict the optimal repair sequence to maximize small, but precise edits? To investigate this, we chose to use ssODN templates to obtain gene edits exploiting single-strand templated repair (SSTR) via the Fanconi anaemia (FA) DNA repair pathway^43^. Previously, numerous studies have focused on enhancing gene editing efficiency using HDR by adjusting the lengths of repair arms^44^, chemically modifying repair templates^45^, or inhibiting DNA repair regulators^46^. Currently, there is no tool to design ssODN repair templates that can predict expected gene editing efficiencies, or that forecasts the expected ratio of intended versus unintended editing outcomes. Thus, we investigated if deep learning based predictive algorithms can fulfil this need.

We used an eGFP to eBFP conversion assay^47^, which depends on the change of two nucleotides (<u>C</u>C<u>T</u> to <u>G</u>C<u>C</u>) to explore the design space by InDelphi predictions. We computed the predicted percentage of on-target repair as a function of both the left and right repair arm lengths and calculated the chance for overall perfect repair as the joint probability of perfect repair occurring between the genome and both repair arms, left and right of the cut (Fig. 5a). Because this extended the use of the InDelphi model beyond previous applications, we termed this approach Pythia, in reference to the priestess at the Greek temple of Delphi in antiquity^48^. We introduce a bioinformatics-based solution for generating what we termed ‘Pythia matrices’ (Fig. 5a, Fig. S10). These matrices depict the predicted gene editing efficiencies in relation to the lengths of both the left and right repair arms. Next, we investigated whether Pythia predictions correlated to experimental observations by designing repair templates for eGFP to eBFP conversion with high and low Pythia scores (Fig. 5b).

**Fig. 5.**
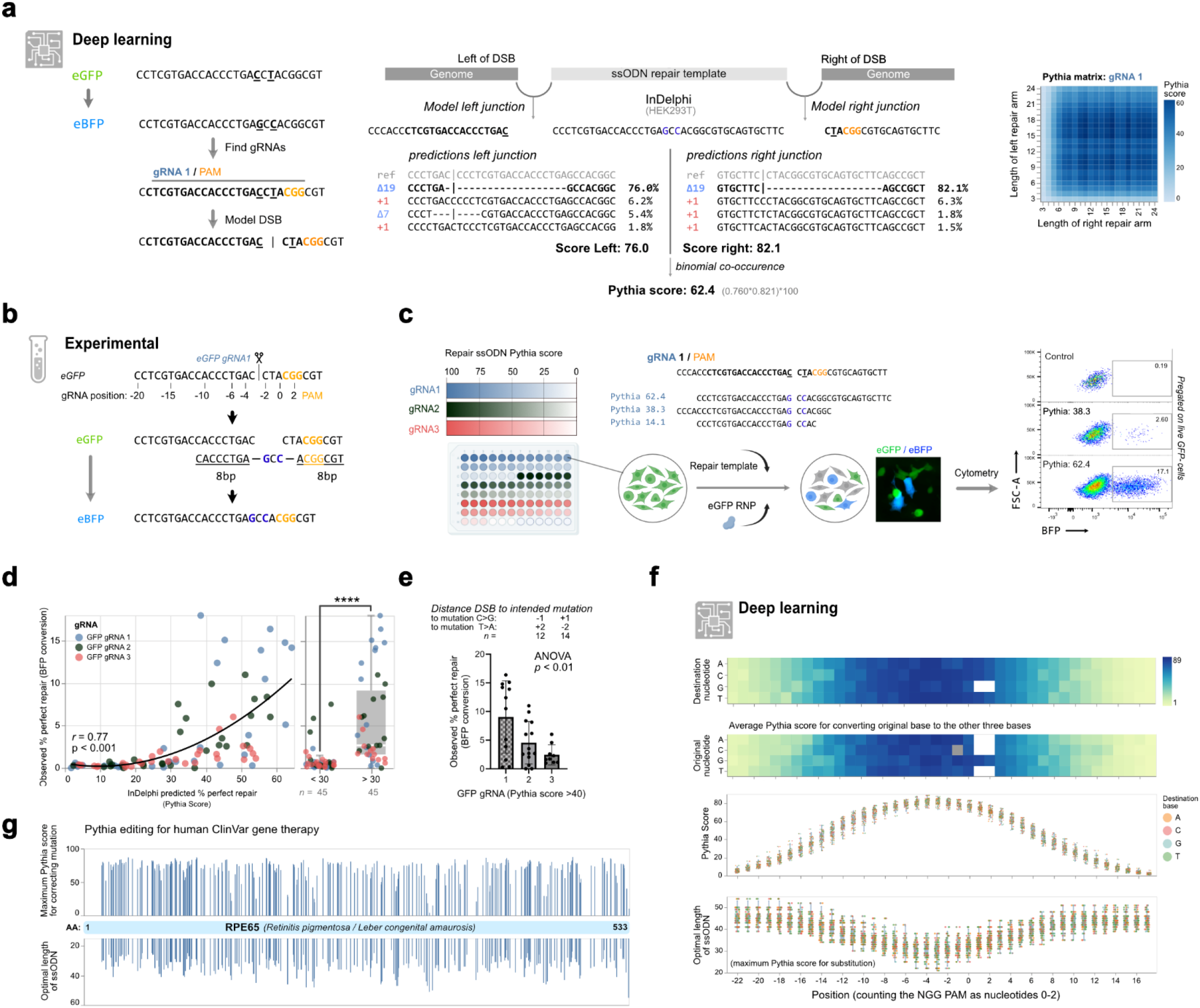
Pythia Editing, leveraging predictability to create small point mutations. **(a)** eGFP-to eBFP conversion can be achieved by establishing two point mutations. Schematic representation of Pythia, an bio-informatics pipeline, deploying the InDelphi model to calculate expected editing outcomes on both junctions, yielding a combined Pythia score defined as the binomial co-occurrence of the intended edit. The Pythia scores for different repair arm lengths is depicted as a Pythia matrix shown on the right. **(b)** Strategy for converting eGFP into eBFP using a 19 bp long ssODN designed by Pythia (homologous sequences underlined). **(c)** Experimental setup for determining eGFP to eBFP conversion efficiencies using three different gRNAs, with 30 distinct ssODN repair templates binned across deciles of Pythia scores. **(d)** Scatter plot showing a direct correlation (r=0.77, p<0.001) between Pythia scores and fluorescence conversion, across all three tested gRNAs. Comparison of conversion rates between ssODN repair templates with a predicted Pythia score of below and above thirty (t-test, p<0.001, n=45). **(e)** The distance between the induced DSB and the site of the intended point mutation influences the median percentage of gene conversion (ANOVA, p<0.01, n=12, 14, 9). **(f)** Modeling of potential Pythia editing outcomes for 35 gRNAs targeting the *X. tropicalis tyr* gene. (Top-to-bottom) Average Pythia score for converting a base to one of the other three bases, plotted first for each destination nucleotide at each position and below for each original nucleotide at each position. Scatter plot of maximum Pythia scores for optimal ssODN design at each position, and the length of optimal ssODN. **(g)** At-scale modelling of Pythia editing for restoring human *RPE65* pathogenic missense variants annotated in ClinVar to restore the wild-type amino acid. For each variant and the closest gRNA, the maximal achievable Pythia score (top) and the length of the optimal repair ssODN repair template (bottom) is shown.

For each eGFP gRNA (*n=3)*, we generated 30 repair templates with three repair templates in each decile of Pythia scores (bin) and quantified eGFP to eBFP conversion rates (Fig. 5c). This revealed gene editing efficiencies of up to 18%, adhering to a monotonic correlation between Pythia prediction matrices and experimentally determined conversion rates (combined Spearman correlation *r=0*.*77*, p<0.001) (Fig. 5c-d, Fig. S11). As the distance from the intended base pair modification to the DSB increased, gene conversion efficiency decreased, a trend accurately predicted by the Pythia matrices (Fig. 5e).

Next, we explored if the predictability would allow us to model the editing window for small, but precise nucleotide substitutions. We computed the maximum Pythia score to establish individual base changes at positions -22 to +17 to all three possible nucleotide substitutions for 35 distinct gRNAs. This revealed no preference in substitution efficiency depending on which nucleotide was at a given position, suggesting that all possible substitutions are theoretically achievable (Fig. 5f). Further, the greater the distance between the intended substitution and the DSB, the lower the highest possible Pythia score was. Thus, a longer ssODN template is needed to achieve an optimal Pythia score (Fig. 5f). Notably, at the individual gRNA level, sequence contexts exerted a profound influence, which led to considerable variation in the optimal ssODN repair length. For instance, for introducing a substitution at position -6, the model predicted an on average optimal length of 32 nucleotides for the ssODN repair template. In summary, the spread in optimal length of ssODN repair templates ranged from 25 to 42 bps and modeling suggested that a window of -11 (mean Pythia score 61.1 ± 6.3) to +4 (mean Pythia score 61.5 ± 4.6) from the cut-site constituted a suitable window for editing. We predicted the optimal ssODN repair templates for gene correction of all *RPE65* missense mutations associated with retinal degeneration and annotated in ClinVar^49^ where a suitable gRNA was found in proximity. We observed an average Pythia score of 84.4±12.5%, with an ssODN length of 33.9±5.7 nucleotides (Fig. 5g). Given that 81% (n=293) of *RPE65* missense mutations could be edited with a Pythia score of >60, we believe Pythia editing could hold promise for clinical applications.

Finally, we questioned whether Pythia editing could be validated by *in vivo* experiments in *Xenopus* embryos. For this, we chose to design a ssODN repair template to introduce two silent mutations (spaced 5 bp apart) in tyrosinase, a gene essential for pigmentation. Pythia predicted highly efficient repair between the right template arm and the genome (17 bp distance to cut site, Pythia score 94), but suboptimal welding between the left repair arm and the genome (24 bp distance to cut site, Pythia score 46), yielding a total Pythia score of 43 (Fig. 6a). In 65% (n=20) of pigmented animals (Class 3, Fig. 6b-d) that received a high dose of the RNP/ssODN mixture, we observed restriction enzyme patterns indicating successful insertion of the desired point mutations (Fig. 6e).

**Fig. 6.**
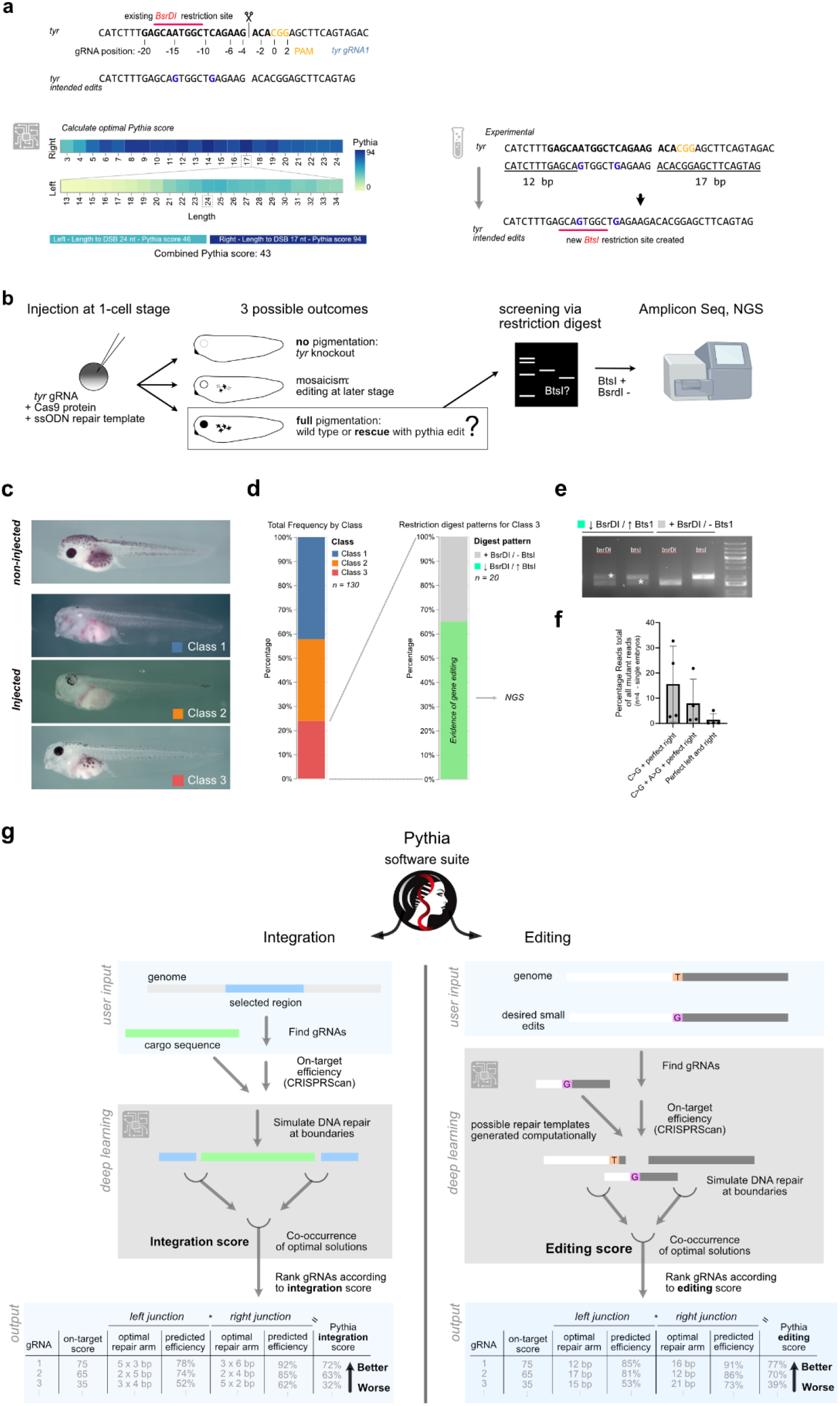
Pythia Editing *in vivo* and Pythia software suite. **(a)** Strategy for establishing two silent point mutations in the *X. tropicalis tyrosinase* gene, using an RNP and a 41 bp ssODN repair template as designed using the Pythia pipeline. **(b)** Schematic of experimental design to detect and quantify successful editing events. **(c)** Compared to non-injected animals, injected animals showed different levels of residual pigmentation, which was binned across three classes (d-e) Frequency of phenotype occurrence grouped by class and evidence for gene editing in class 3 animals according to restriction enzyme analysis.>60% of animals showed evidence of gene editing. **(f)** Quantification of NGS amplicon read analysis. **(g)** Schematic of the Pythia software suite and algorithms for designing integration or editing strategies. The software is freely accessible at pythia-editing.org.

Interestingly, NGS in four animals with pronounced changes in restriction enzyme digest patterns, revealed efficient repair between the ssODN and the genome right of the DSB that also included the C>G edit at 16%+-7.5% of the reads (Fig. 6f). Of those reads of 51% also contained the expected A>G editing (8%+-4.8% of total reads). However, as predicted by Pythia, repair between the left repair arm and the genome was less efficient (24 bp, Pythia score 46) and only 1.4%+-1.42% of the reads contained perfect repair on the left and the right arm, with the establishment of the two silent point mutations.

Together, this provides first proof-of-principle demonstration of SSTR-mediated single base pair substitutions *in vivo* with ssODN repair templates rationally designed via deep learning approaches.

We provide a freely accessible web tool (pythia-editing.org) to allow custom design strategies for base edits or integration using Pythia (Fig. 6g). As the predictive power of the Pythia suite can be extended to calculate optimal overhang arms for integration events, we are able to provide more liberal options that extend beyond the trimology design and may provide even higher integration efficiencies, protection of junction boundaries and in-frame gene-tagging.

## Discussion

Improvements in gene editing strategies often rely on rational design or systematic protein engineering^50^. In contrast, pre-trained deep learning networks are considered enigmatic entities, often referred to as “black boxes”^51^. Nonetheless, deep learning consolidates vast amounts of data from complex biological experiments into models, which can be parsed to gain insights into principles involved and guide experiments to confirm new hypotheses parsed from predictive models^52^.

Here, we utilized a pretrained model (InDelphi^17^) towards optimizing both large cargo integration (trimology integration) and gene editing (Pythia). Exploiting a system of microhomologies tandem repeats, our key finding is that DNA repair is predictable at the interface between Cas9-mediated breaks and exogenously delivered DNA, both *in vitro* as *in vivo*. We distil a rule set for selecting gRNAs driving high integration efficiencies and for designing said target-specific repair templates. As such, rational design of donor repair arms to maximize desired editing outcomes is achievable, substantially aided by the deep learning network, and delivered mechanistic insights into how genomic context (G at nucleotide position -4) impacts the efficiencies of gene integration. Interestingly, this is inverse to the relationship between the nucleotide at position -4 and the propensity to repair via a +1 insertion^17,19^. Of note, the presence of a G at this position predicts blunt DSB induction by CRISPR/Cas9^53^, possibly directly connecting the Cas9 incision type to the preferential engagement of distinct DNA repair pathways. Together this allows for gRNA selection and rational design of repair arms via deep learning approaches on a sequence context-specific manner.

These findings have numerous applications in biotechnology. Some of the advantages of trimology integration are that it is directional, single copy, and locus specific, thus avoiding most of the drawbacks of other *in vivo* transgenesis techniques, such as positional effects in enhancer screenings^37,38^. We have demonstrated that trimology integration enables cargo insertion in human and *Xenopus* and facilitates endogenous tagging *in vivo* of mouse brains. A key advantage of trimology repair arms are their exceptionally short lengths (6-15 bp), which simplifies the generation of repair templates that can be efficiently produced using straightforward overhang PCR methods, facilitating large-scale cell screening projects.

By co-opting the MMEJ repair pathway, trimology integration is applicable in certain cellular contexts when homology-directed repair (HDR) is known to be inefficient or even inactive, such as in early developing vertebrates^44^ or post-mitotic adult tissues such as retina or brain^54^, providing potential for gene therapy approaches^55^. Moreover, its applicability extends to CAR-T manufacturing^56^, where the integration of protective base pairs, deemed necessary to counteract deletions into the genome or cassette, could be replaced by trimology arms to ensure predictable repair^57^.

Together, we show that small trimology integration with tandem repeat repair arms is sufficient for predictable in-frame repair and offers higher predictability then NHEJ-based methods^30,58,59^, which are error-prone. Importantly, we demonstrate that trimology repair arms safeguard the genome and the donor template during DNA integration, precluding extensive genomic deletions.

Next, we demonstrate the potential for transfer learning of pretrained deep learning models (such as InDelphi) towards optimizing gene editing. We establish a metric called Pythia score that provides a predictive measure towards the efficiency of establishing intended point mutations, but not bystander mutations, used CRISPR-mediated SSTR with ssODN repair templates. As such, rational design of mass-producible small ssODN repair templates specifically designed to maximize gene editing is possible.

We also present the first proof-of-principle demonstration for single base pair substitutions in rapidly developing *Xenopus* vertebrate embryos with simple ssODN repair templates, without resorting to host transfer methods^60^. These templates can be strategically designed with the help of the Pythia design tool. Although our efficiency rates are modest, they align with previous studies conducted in rapidly developing zebrafish embryos^44,61^. Importantly, there’s potential to enhance these efficiencies through modifications to the ssODNs^45^ or addition of small interfering molecules targeting mediators of DNA repair pathways^62^. Next, these findings suggest that DNA repair outcomes can also be predictably influenced when using ssODN templates. This opens up new possibilities for enhancing Pythia-based integration by using ssODN or hybrid ssDNA templates to further reduce off-target integration and cellular toxicity^63^.

One limitation of our methodologies is their dependence on double-strand breaks (DSBs), which are known to activate the p53 pathway and can sometimes result in complex genomic rearrangements, including genomic deletions, chromosomal translocations, and chromothripsis. While several alternative strategies, such as base editing^64^, prime editing^26^, and integrase-based approaches^27–29^, offer potential solutions, they are not without their own challenges. Meanwhile, base editing is constrained by its editing window, limited in the variety of genetic substitutions it can achieve^64^. Nonetheless the limitations associated with DSBs, this is to our knowledge the first description of using sequence-context specificity to optimize gene integration and gene editing, thus showing that there is at least some determinism in DNA repair and opening avenues for further optimizing any current or future gene editing tool that relies on DSB repair.

Indeed, deep learning (DL) has been shown to predict outcomes for CRISPR/Cas9^17^, base^65^ and prime editing^66^ effectively. In our study, we reveals for the first time an unanticipated level of non-randomness of DNA repair on the interface between the genome and exogenous donor DNA, which is explainable by deep learning models trained on CRISPR/Cas9-induced DSB repair^17^. While this transfer learning approach is validated by our experiments, specific fine-tuning of such models on cell specific repair patterns would further increase the reliability of predictive models. We believe that our findings open an unexplored design space using deep learning to optimize genome rewriting and will serve as a primer for training additional cell-type specific models to further optimize genome editing^67^. This may have profound implications for clinical *in vivo* CRISPR/Cas9-mediated gene therapy approaches.

To facilitate easy access, we have created an online tool for automated design of repair templates for both trimology integration as well as for Pythia, to allow straightforward and user-friendly access to these powerful technologies (pythia-editing.org). Drawing inspiration from the ancient world, we have named our approach “Pythia” after the high priestess at the Temple of Apollo in Delphi. Renowned for her perceived ability to foretell the future, the Pythia was a revered figure whose prophecies guided countless decisions in antiquity^48^. Like Pythia, our methodology predicts and influences outcomes, albeit in the realm of CRISPR/Cas9 genome editing.

## Material And Methods

### Cell culture

HEK293T (ATCC, CRL-11268) were cultured as recommended by the ATCC. Cell lines tested negative for mycoplasma and were authenticated by the suppliers.

### Modelling of gene editing outcomes

The InDelphi-model was obtained from https://github.com/maxwshen/inDelphi-model and deployed in a suitable Python virtual environment (see code availability). To investigate the impact of the number of tandem repeats of trimologies on the expected percentage of perfect DNA repair, we developed custom Python code. % repair via µH is defined as the sum of all repair outcomes that use at least one trimology. The code iteratively analyzes trimology lengths ranging from two to six and the number of tandem repeats from one to eight. This analysis was conducted using either the InDelphi-HEK293T or InDelphi-mESC predictive model for the first 250,000 gRNA sites identified by presence of an NGG PAM, encountered in the human gencode v43 transcript sequences.

### Cloning and *in vitro* linearization of PaqMan repair plasmids and PCR-generation of repair templates

Donor plasmid was assembled in a pUC19 backbone using Gibson cloning (NEBuilder® HiFi DNA Assembly Master Mix) and feature a pCMV-eGFP transgenic cassette flanked by either zero, 4x or 5x trimology arms and inverted PaqCI restriction enzyme sites. Insert was obtained from AAV-CMV-GFP which was a gift from Connie Cepko (Addgene plasmid # 67634 ; RRID:Addgene_67634). pUC19 destination vector was commercially purchased (N3041S, NEB). Inverted PaqCI sites and trimology repair arms were added by overhang PCR prior to Gibson assembly. Linearization was performed by overnight digest at 37C° of 10 µg of donor plasmid using 20 Units of PaqCI (R0745, NEB) in 1x rCutSmart buffer (B6004S, NEB). Complete linearization was ensured via classical agarose gel electrophoresis.

Alternatively, repair templates containing trimology repair arms were generated by overhang PCR approaches using Phusion polymerase (Thermo-Scientific) with primers designed to contain an overhang sequence containing the trimology repair arms (listed in Supplementary Table 3). For *in vitro* use, PCR products were cleaned via MinElute PCR Purification Kit (28004, Qiagen) and eluted in ultrapure water. For *in vivo* use, PCR products were cleaned via classical phenol/chloroform extraction with Sodium Acetate / EtOH precipitation and quantified using Nanodrop (Thermo-Scientific).

### Trimology integration *in vitro*

*Aavs1* gRNA was assembled using Alt-R® CRISPR-Cas9 IDT crRNA and Alt-R® CRISPR-Cas9 tracrRNA, according to the manufacturer’s instructions, by heating it up to 95°C and cooling it down to RT yielding a duplex at a final concentration of 1 μM. Cas9 protein (PNABio, CP01) was diluted to 166.67 ng/μL in 1x PBS. HEK cells where reverse transfected using Lipofectamine CRISPRMAX (Thermo-Scientific) as follows. RNP was assembled via incubation for 5 minutes at room temperature of 1 μM gRNA duplex, 250ng of Cas9 protein and 0.6 μL Cas9 PLUS reagent (from CRISPRMAX kit) in 23μL Opti-MEM (Thermo-Scientific). After, 200 ng of PaqCI digest product was added to the RNP. Transfection complexes were generated by incubation at room temperature for 20 minutes of 25 μL of RNP/repair template, 1.2μL of CRISPRMAX transfection reagent and 23.8 μL of Opti-MEM medium. Resulting transfection complexes were mixed with 40,000 HEK293T cells (suspended in a total volume of 100 μL of DMEM) and plated on 96-well Nunclon plates (Thermo-Fisher). Cells were cultured for 25 days and cell sorting for GFP+ cells was performed.

For 32-target experiment, gRNAs were designed for CDS from human genome assembly GRCh38 using a custom python script, identifying gRNAs with each permutation of strong (S) and weak (W) bases at position -6 to -4, and AGG at position -3 to -1 with NGG as PAM at position 0 to 2. Identified gRNAs where filtered for those with CRISPRScan scores^68^ exceeding 80. In order to avoid negative selection due to gene essentiality when targeting CDS, we filtered the gRNA list to exclude any gene occurring in the DEG15^31^, a database of essential genes as determined from shRNA and CRISPR/Cas9 screens. Next, the eight classes of permutations involving S and W bases were sorted into bins. For each class, one guide RNA (gRNA) was selected per bin, arranged according to the degree of sequence context microhomology, ranging from low to high. For each gRNA off-target profile was determined, and deemed acceptable, via Cas-OFFinder^69^ (list of gRNAs in Supplementary Table 2).

Gene editing was performed identical to above, with following exceptions. We used 100ng of repair template (instead of 200 ng), generated by overhang PCR as described above from AAV-CMV-GFP (Addgene plasmid # 67634; RRID:Addgene_67634) (Supplementary Table 3). Transfection was performed one-day after seeding of 25,000 cells in a 96-well plate. Cells were sorted at day 2 and at day 15. Here, integration efficiencies is defined as follows: all cells pre-gated on live cells, via SYTOX™ Deep Red Nucleic Acid Stain (1 µM final) (Thermo-Scientfic). Then, (%GFP+ cells at day 15*(100/%GFP+ cells at day 2), thus accounting for differences in initial transfection efficiency via transient expression of the pCMV:eGFP cassette at Day 2. On-target efficiency is defined as follows: ((Integration efficiency on-target gRNA) – (Integration efficiency neg. control gRNA)) at day 15, thus identifying the level of true on-target integration.

### Trimology integration *in vivo* in *Xenopus*

*Xenopus* animals were kept according to the Swiss law for care and handling of research animals. Husbandry and treatment were approved by the local authorities (Veterinäramt Zürich). Gene symbols follow Xenbase (http://www.xenbase.org/, RRID:SCR_003280). For *Xenopus* experiments, repair templates for pCMV:eGFP, Pax8-CNS1:eGFP and CarAct:dsRed2 were generated by overhang PCR as described above. For pCMV:eGFP (5x trimology) and Pax8-CNS1:eGFP experiments, *Hipp11*-α and *hipp11*-β gRNAs were assembled as follows: 1 µL Alt-R® CRISPR-Cas9 IDT crRNA (100 µM stock) and 1µL Alt-R® CRISPR-Cas9 tracrRNA (100 µM stock) were mixed with 3 µL of Nuclease-Free Duplex Buffer (IDT), and heated at 95C° for 5 minutes, and allowed to cool to room temperature. The following mix was made for RNP assembly: 1.8 μL of Cas9 protein (1 μg/μL, PNABio CP01) with 0.2 ul of gRNA, heated up to 37°C for 5 minutes, before adding repair template. The final injection mix consisted of 1 μL of *hipp11*-α RNP, 1 μL of *hipp11*-β RNP and 1 μL of repair template (stock concentration 10ng/ul) thus yielding a final repair template concentration of 3.33 pg/nL. Embryos were injected unilaterally in the two-cell stage. For pCMV:eGFP (4x trimology) and CardiacActin:dsRed2, we mixed Cas9 protein (3 μL at 1 μg/μL, PNABio CP01), with gRNA (1 μL) and incubated for 5 minutes at 37°C to assemble RNP. RNP was mixed with repair template at a ratio of 4:1, thus adding 1 μL of repair template (10 ng/ul) to the mix yielding a final repair template concentration of 2 pg/nL. Embryos were injected in the one-cell stage immediately after cortical rotation, targeting the grey sperm entry point with 5-10 nL of injection mix.

Embryo development was monitored and at Nieuwkoop-Faber stage 40, embryos were lysed (50 mM Tris pH 8.8, 1 mM EDTA, 0.5% Tween-20, 2 mg/ml Proteinase K) overnight at 55°C. Three classes of embryos were lysed as follows: Embryos with unilateral or bilateral non-mosaic fluorophore expression, embryos with mosaic expression often restricted to a subset of the muscle cells and control embryos of the same clutch which were not micro-injected. After Proteinase K inactivation, junction products between the *h11* locus and transgene cassette were picked up using PCR and subjected to Sanger sequencing. Whole-embryo bleaching, staining and clearing was performed as previously described^70^, using here 1:250 anti-GFP (Aves, GFP-1020) and 1:250 anti-RFP (Rockland, 600-401-379-RTU, crossreacts with mCherry).

### Trimology integration *in vivo* in mouse brain

pAAV-mTubb3 and pAAV-nEFCas9 was a gift from Juan Belmonte (Addgene plasmid #87116; RRID:Addgene_87116, Addgene plasmid # 87115; RRID:Addgene_87115). Sequence between AgeI and NdeI restriction sites was exchanged for a synthetic DNA fragment containing a gRNA targeting mTubb2a (5’-GGGCGAGTTCGAGGAGGAGG-3’), 3’ Tubb2a sequence context, Trimology arms and eGFP to generate pAAV-mTubb2a. pAAV-nEFCas9 and pAAV-mTubb2a were packaged with serotypes 8 and were generated by Viral Vector Facility at the University of Zurich.

All procedures of mouse animal experimentation were carried out according to the guidelines of the Veterinary Office of Switzerland and following approval by the Cantonal Veterinary Office in Zurich (licenses 008/2022). Four C57BL/6 mice were used for virus injections. Mice were anesthetized with 1.5-2% isoflurane mixed with oxygen, and were head-fixed in a stereotactic frame (Kopf Instruments). Body temperature was maintained at ∼37°C using a heating pad with a rectal thermal probe. Vitamin A cream (Bausch & Lomb) were applied over the eyes to avoid dry eyes. After analgesia treatment (extended buprenorphine release EthiqaXR, 3.25 mg/kg, subcutaneous; lidocaine over scalp), an incision was made on the scalp, and small holes were drilled over bilateral visual cortex using the following coordinates: 3.5 mm caudal, 2.5 mm lateral relative to bregma, 0.5 mm ventral from the pia. We used 1:1 mixture of AAV-Cas9 (1.5 × 10^13^ GC ml^−1^) and AAV-mTubb3 (2.3 × 10^13^ GC ml^−1^), and injected 600 nl of AAVs in each hemisphere. To prevent virus backflow, the pipette was left in the brain for 5–10 min after completion of injection. Mice were housed for three weeks to allow for gene knock-in. After, animals were euthanized, perfused using 4% PFA, brains dissected and post-fixed for 2 hours in 4% PFA. Whole-brain staining was performed as follows and adapted from previously described WildDisco^42,71^. Whole-mount brains where dehydrated to 100% MeOH, bleached in and delipidated with Dichloromethane. Next, brains were permeabilized/blocked for 3 days using 10% donkey serum and 2% Triton X-100 in 0.1 M PBS. Antibody staining was performed with a 1:250 anti-GFP (Aves, GFP-1020) and 1:250 anti-RFP (Rockland, 600-401-379-RTU) in 5mL immunostaining buffer containing 3% Donkey serum, 10% CHAPS, 2% Triton X-100, 10% dimethylsulfoxide (DMSO), 1% glycine, 1% CD5 in 0.1 M PBS for 7 days at 37C° on a rotating wheel. After 3 days of washing, 1:400 of donkey-anti-rabbit-Cy3 antibody (Jackson ImmunoResearch, 711-165-152) and donkey-anti-chicken-AlexaFluor594 antibody (Jackson Immuno Research, 703-585-155) in immunostaining buffer was added for 7 days at 37C° on a rotating wheel. Brains where washed extensively, dehydrated to 100% MeOH in steps and then cleared in BABB.

### Imaging Methods

For stereomicroscopy imaging, a SteREO Discovery.V8 from Zeiss and Zen2011 Blue Edition was used. *In toto* cleared *X. tropicalis* embryos and mouse brains were imaged using mesoSPIM^71,72^. For all mesoSPIM recordings, fluorophores were excited with the appropriate laser lines and a quadband emission filter (ZET405/488/561/640, Chroma) was employed. Imaging was performed using dibenzyl ether (DBE) as immersion medium. Two-photon imaging was performed with a reflective multi-immersion Schmidt objective as described before^73^. Live time-lapse imaging for *Xenopus* embryos was performed on a Widefield Thunder Imager (Leica). Stitching was performed with BigStitcher^74^. Data was rendered using Imaris (Oxford Instruments) or Napari (https://github.com/napari/napari)^75^. U-net mediated segmentation was performed as described before^70^.

### DNA preparation, Sanger and next-generation sequencing

Cells, *Xenopus* embryos or mouse AAV-injected hemispheres were lysed (50 mM Tris pH 8.8, 1 mM EDTA, 0.5% Tween-20, 2 mg/ml Proteinase K) at 55°C overnight. After Proteinase K inactivation (10 minutes incubation at 98C°), PCRs were performed using GoTaq G2 (Promega), Q5 (NEB) or Phusion polymerase (Thermo-Scientific) (primers listed in Supplementary Table 1). For sequencing, amplicons were cleaned using nucleoSpin Gel and PCR Clean-up (Machery-Nagel) and sent for commercial sequencing (Microsynth, Balgach, CH). For NGS, amplicons were generated by PCR with appropriate adapter sequences and commercially sequenced (INVIEW CRISPR Check (size: 450 - 500 bp), Eurofins Genomics). Data analysis was performed using CRISPResso2^76^ and/or custom data-processing.

### Pythia *in silico* modeling

The Pythia Python script is designed to simulate CRISPR/Cas9-mediated gene editing efficiencies using a given wild-type (WT) and mutant DNA sequence. It iteratively constructs potential editing templates by varying the lengths of the left and right homology arms and utilizes the inDelphi tool to predict repair outcomes and their frequencies. The results, including the predicted repair outcomes and their corresponding frequencies, are stored and reported, in order to identify the most effective repair template for achieving the desired genomic modification.

We modelled the optimal ssODN repair template length, with the maximal Pythia score, across clinically relevant point mutations in RPE65, involved in -among others-Retinitis pigmentosa and Leber congenital amaurosis. For this, we obtained all RPE65 ClinVar (accessed at 06/01/2024) single nucleotide missense variants. For each missense variant, we calculated the minimal amount of base changes required to change the codon usage from the human missense variant amino acid towards the restoration of the wild-type amino-acid at that location. Next, Pythia code was employed to compute the optimal ssODN repair template with the maximal Pythia score to establish this base point mutation, thus reverting the clinically relevant mutation at the amino acid level.

### Pythia editing *in vitro*

Potential ssODN repair templates were designed for three independent GFP gRNAs in order to establish two point mutations to convert eGFP to eBFP. Pythia scores were calculated with repair arm length set at 1 to 24, both left and right. From these we performed a binning from 0-100 across the scores and randomly selected 30 repair templates for each gRNA, selecting three repair templates per decile bin, thus 90 total. ssODN repair templates were ordered as desalted non-modified primers from Microsynth AG (Balgach, Switzerland) (Supplementary Table 4). HEK293T-*AAVS1*(CMV:eGFP), featuring a stable one-copy integration of a pCMV:eGFP construct, were seeded at a density of 10,000 cells in a 96-well plate in 150μL of standard DMEM. 24 hours later, cells were transfected using Lipofectamine CRISPRMAX (Thermo-Scientific) and Lipofectamine 3000 (Thermo-Scientific) as follows. gRNA was assembled using Alt-R® CRISPR-Cas9 IDT crRNA and Alt-R® CRISPR-Cas9 tracrRNA, according to manufacturer’s instructions, by heating it up to 95°C and cooling it down to RT yielding a duplex at a final concentration of 1 μM. RNP was assembled via incubation for 5 minutes at room temperature of 1 μM gRNA duplex, 250ng of Cas9 protein (Alt-R™ S.p. Cas9 Nuclease V3, IDT) and 0.6 μL Cas9 PLUS reagent (from CRISPRMAX kit). Transfection complexes for RNP were generated by incubation at room temperature for 20 minutes of 25 μL of RNP/repair template, 1.2μL of CRISPRMAX transfection reagent and 23.8 μL of Opti-MEM medium. Transfection complexes for ssODN were generated using lipofectamine 3000 according to the manufacturer’s instructions. In brief, 1ul of 20 nmol/mL of ssODN repair template was packaged in a final volume of 10 μL. Both, RNP transfection (50 µl final per well) and ssODN transfection reagents (10 µl final per well) were added to the 96-well plate. Next day (approximately 20 hours later), medium was refreshed, and cells were split and maintained according to standard HEK293T principles, until analysis via flow cytometry at day 18.

### Pythia editing *in vivo* in *Xenopus*

A gRNA targeting *X. tropicalis tyrosinase* was designed and the Pythia software was deployed to identify the optimal repair template to generate a double point mutation. Repair template was ordered as desalted ssODN from Microsynth AG (Balgach, Switzerland). gRNA was assembled using Alt-R® CRISPR-Cas9 IDT as described above for *Xenopus*. The following mix was made for RNP assembly: 3 μL of Cas9 protein (1 μg/μL, PNABio CP01) with 1 ul of gRNA, and incubated at 37°C for 5 minutes. Next, 1 µL of ssODN repair template (5 µM stock, 1 µM final concentration) was added. Embryos were micro-injected in the one-cell stage immediately after cortical rotation, targeting the grey sperm entry point with 5-10 nL of injection mix. Restriction digests of PCR products where performed with BsrDI (New England Biolabs, R0574S) overnight at 37°C in NEB buffer r2.1 and with BtsI-v2 (New England Biolabs, R0667S) overnight at 37°C in NEB rCutSmart.

## Supporting information

SI Videos 1-5.zip

## Acknowledgements

The authors thank Daniel Invernot Pérez, Marko Vujanovic and Daniel Cardoso Prata for technical assistance and Julia Traversari for *Xenopus* animal care and husbandry. Further, we acknowledge Jean-Charles Paterna and the UZH Viral Vector Facility (www.vvf.uzh.ch) for production of AAV. General histology was performed at the histology core of the Institute of Anatomy, supported by Shunmugam Nagarajan. Imaging was performed with equipment maintained by the Center for Microscopy and Image Analysis, University of Zurich. Flow cytometry was performed with equipment of the flow cytometry facility, University of Zurich. T.N. received funding from H2020 Marie Skłodowska-Curie Actions (xenCAKUT - 891127) and S.S.L. from an ERC Starting Grant (grant agreement no. 804474, DiRECT) by the European Union’s Horizon 2020 Research and Innovation Program. Further funding support came from the Swiss National Science Foundation (310030_189102 to S.S.L; 310030_192617 to F.H.; Ambizione Grant PZ00P3_216312 to S.H); the URPP Adaptive Brain Circuits in Development and Learning (AdaBD) of the University of Zurich (to F.H. and R.B-G.); and the US Brain Initiative (1U01NS090475-01, F.H.).

## Competing Interest Statement

T.N. and S.S.L. have filed a patent application (EP23192134.7) in relationship to this work.

## Code availability

The inDelphi model is available online at https://indelphi.giffordlab.mit.edu/ and at https://github.com/maxwshen/inDelphi-model. The design tools can be accessed via GUI at pythia-editing.org. Further code is available upon request.

## Supplementary information

### Supplementary Figures

**Supplementary Figure 1:**
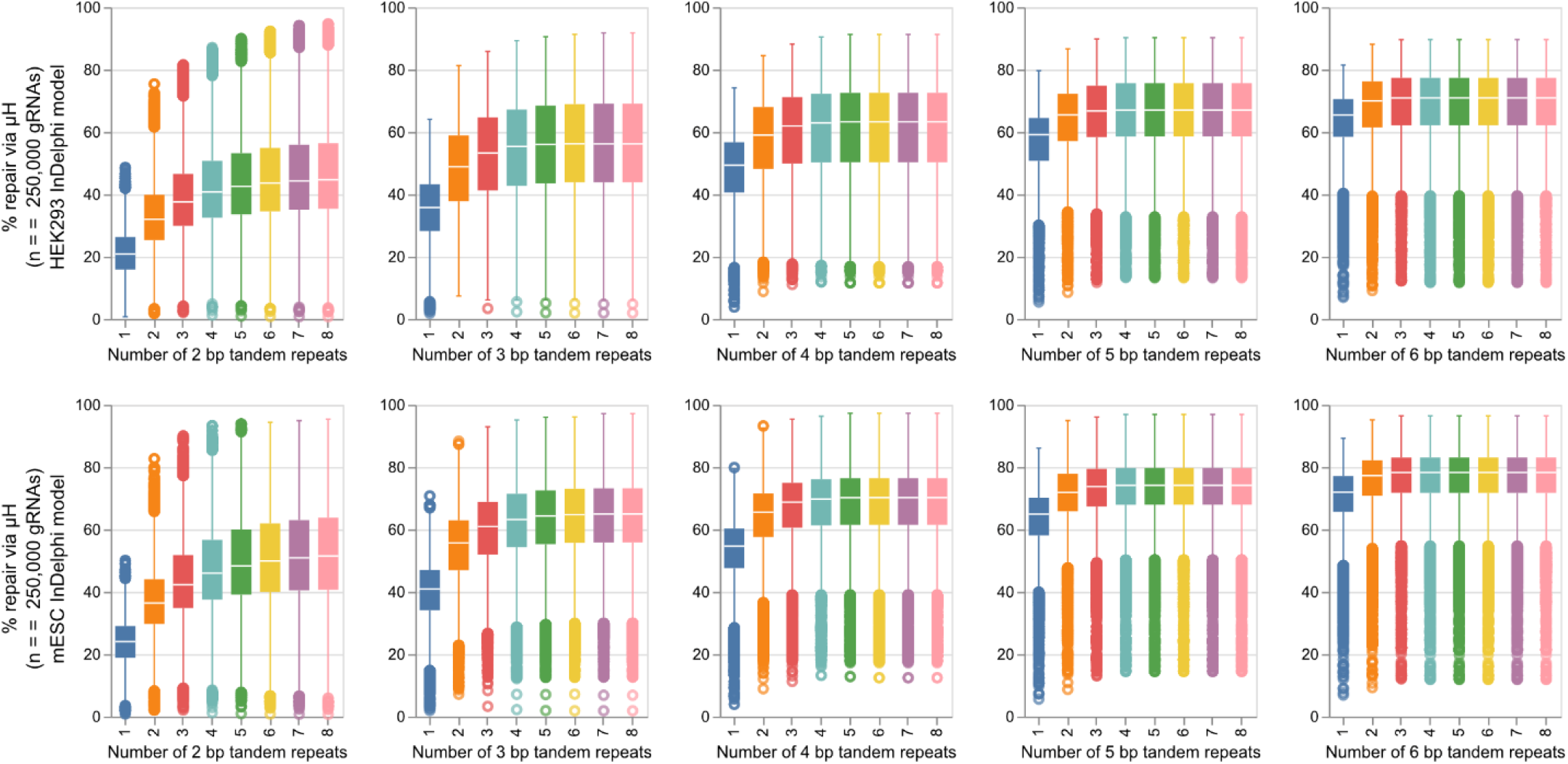
Computational modelling of predicted %repair by µH in relationship to a number and length of tandem repeats. For 250,000 gRNAs binding the human genome, we modelled the expected editing outcomes when adding local sequence context left of the CRISPR/Cas9-mediated DSB to the right of the cut. Predicted %repair by µH is defined as any repair that mobilizes any of the available tandem repeats. This was computed using the InDelphi-HEK293T model (Top) and the InDelphi-mESC model (bottom).

**Supplementary Figure 2:**
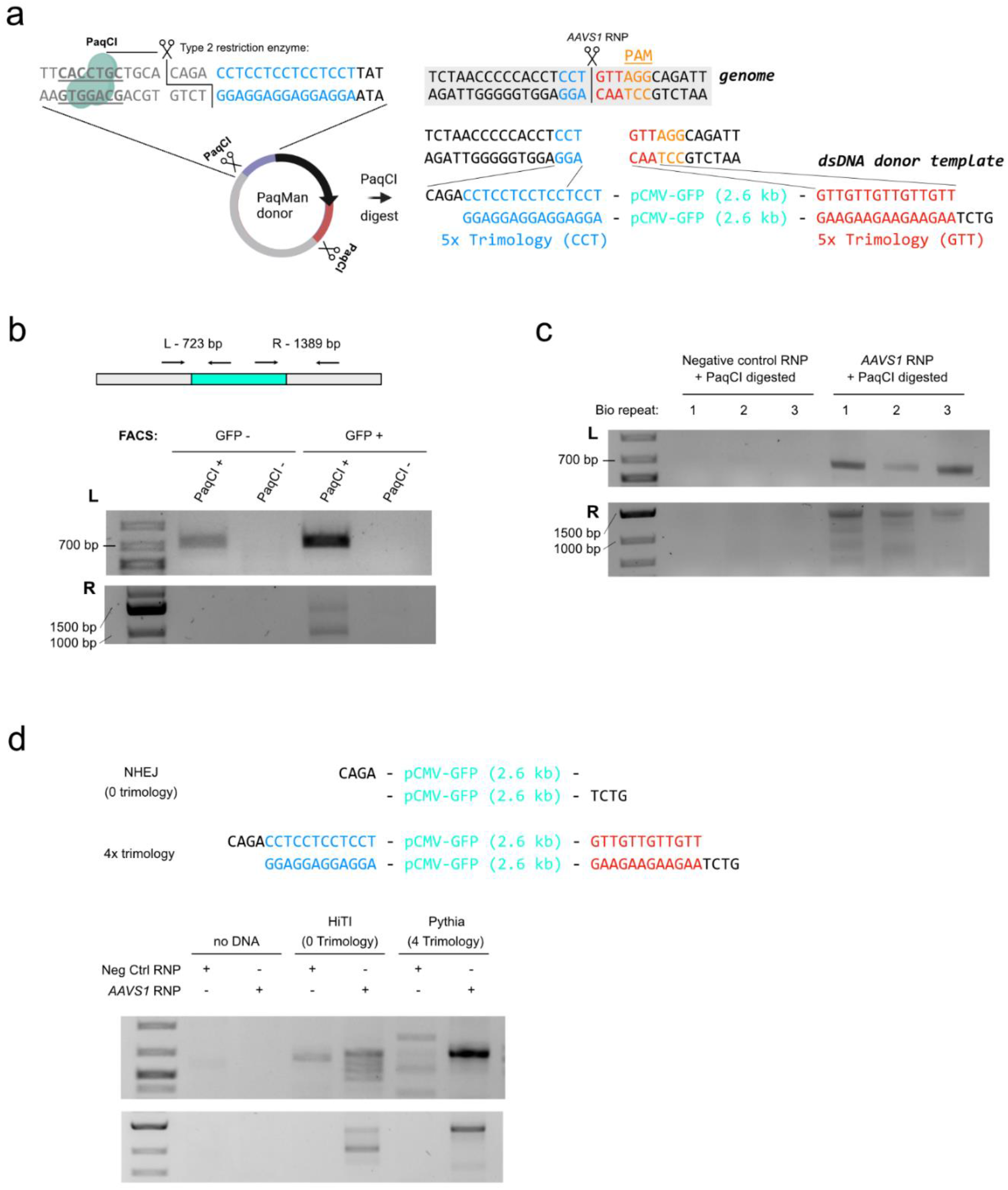
Trimology integration in the *AAVS1* locus. **(a)** Scheme of CRISPR/Cas9 integration strategy. The donor plasmid consists out of pCMV-eGFP expression cassette, flanked by 5x Trimology arms and inverted PaqCI restriction enzyme binding sites. dsDNA donor template is liberated by *in vitro* PaqCI digest, where this type II restriction enzyme provides a cut away from its recognition sequence, allowing fully customizable edges of the dsDNA donor template. This digest is co-delivered with *AAVS1* RNP into HEK293T cells. **(b)** Only with digested (PaqCI-linearized), but not with undigested, can we amplify 5’ (L) and 3’ (R) junction products. This was performed on lysis from cells FACS-sorted into GFP- and GFP+ subpopulations. **(c)** 5’ (L) and 3’ (R) junction products can only be amplified when co-delivering *AAVS1*, and not when co-delivering negative control RNP. **(d)** dsDNA repair templates as obtained after *in vitro* PaqCI digest of donor plasmids. This repair templates where co-delivered together with either negative control RNP or *AAVS1* RNP and contain either 4x trimology repair arms or 0x trimology repair arms (NHEJ).

**Supplementary Figure 3:**
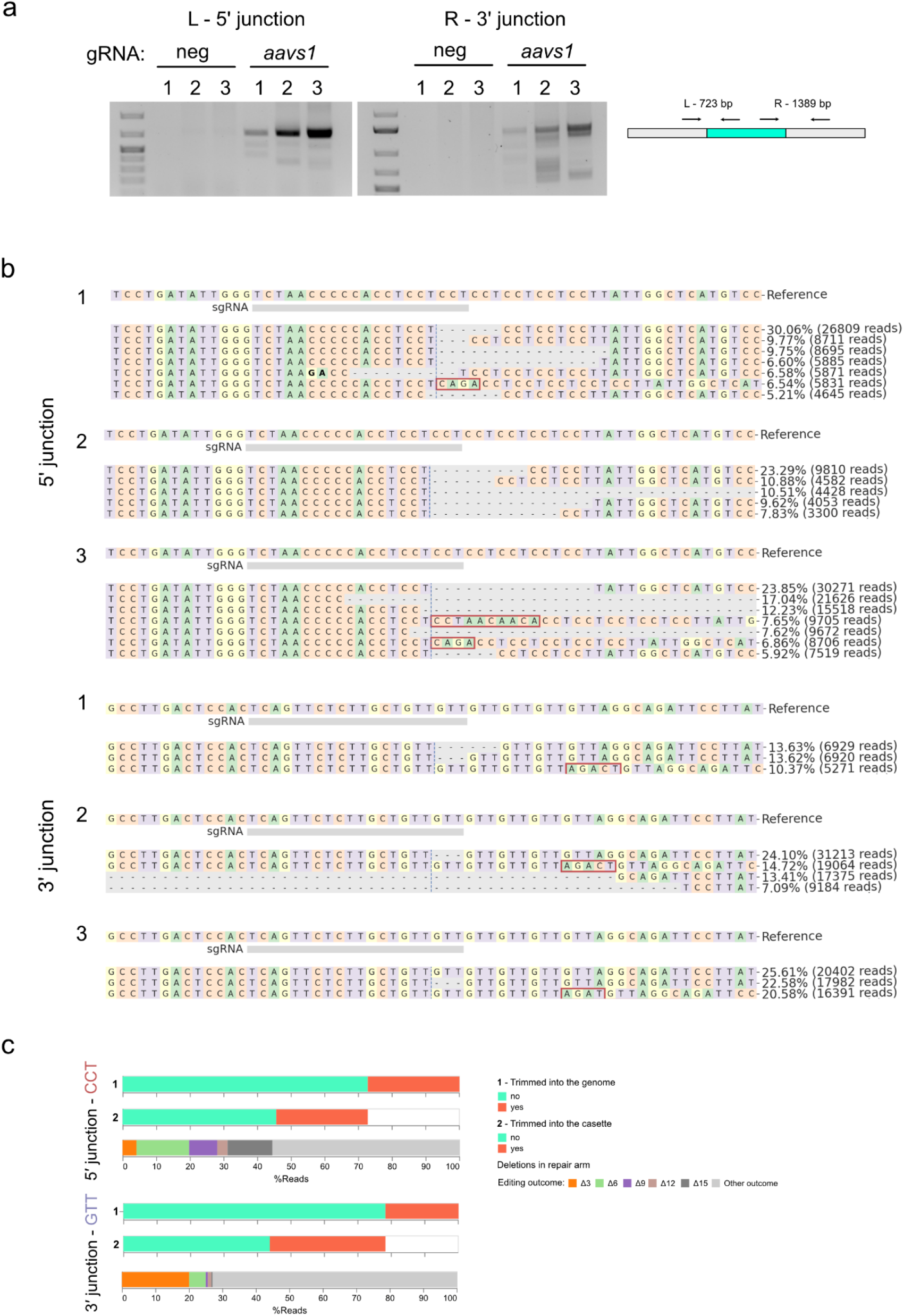
Targeted deep amplicon sequencing of genome-transgene boundary products. **(a)** 5’ (L) and 3’ (R) junction products can only be amplified when co-delivering *AAVS1*, and not when co-delivering negative control RNP. Numbers 1 through 3 denote biological repeats (*n*=3). **(b)** CRISPResso2 analysis of next-generation sequencing of 5’ (L) and 3’ (R) junction products. **(c)** Visualisation of genome editing outcomes on both genome-transgene junctions demonstrating trimming both into the genome (1) and the transgene cassette (2). For each read in which no trimming into the genome or the cassette was observed, the number of deletions in the repair arms are shown.

**Supplementary Figure 4:**
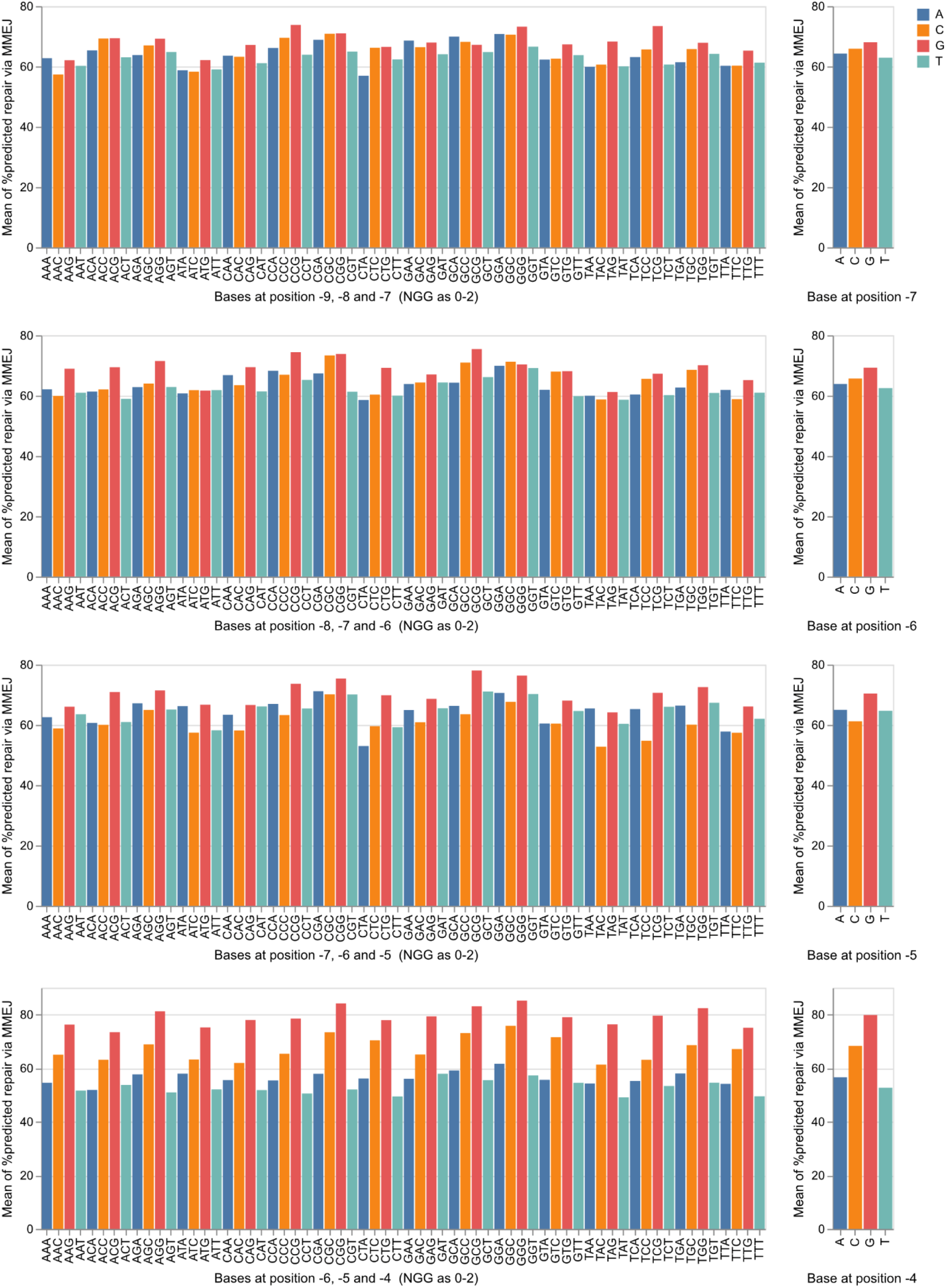
Modelling the predicted percentage of repair via MMEJ based on the base-composition of the gRNA binding site in the genome reveals that base composition at position -4 increases the % of gene editing outcomes by MMEJ for gRNAs with PAM “NGG”. We performed an exome-wide analysis across the human coding genome using the InDelphi-HEK293 model retrieving 10,813,171 unique gRNAs. A subset of 500,00 gRNAs was randomly selected for plotting. This reveals that there is an enrichment for expected % of gene editing outcomes by MMEJ when the base at position -4 is a C or G, when compared to A or T. Such enrichment is not apparent for the positions -5 through -9 (counting the NGG PAM as nucleotides 0-2).

**Supplementary Figure 5:**
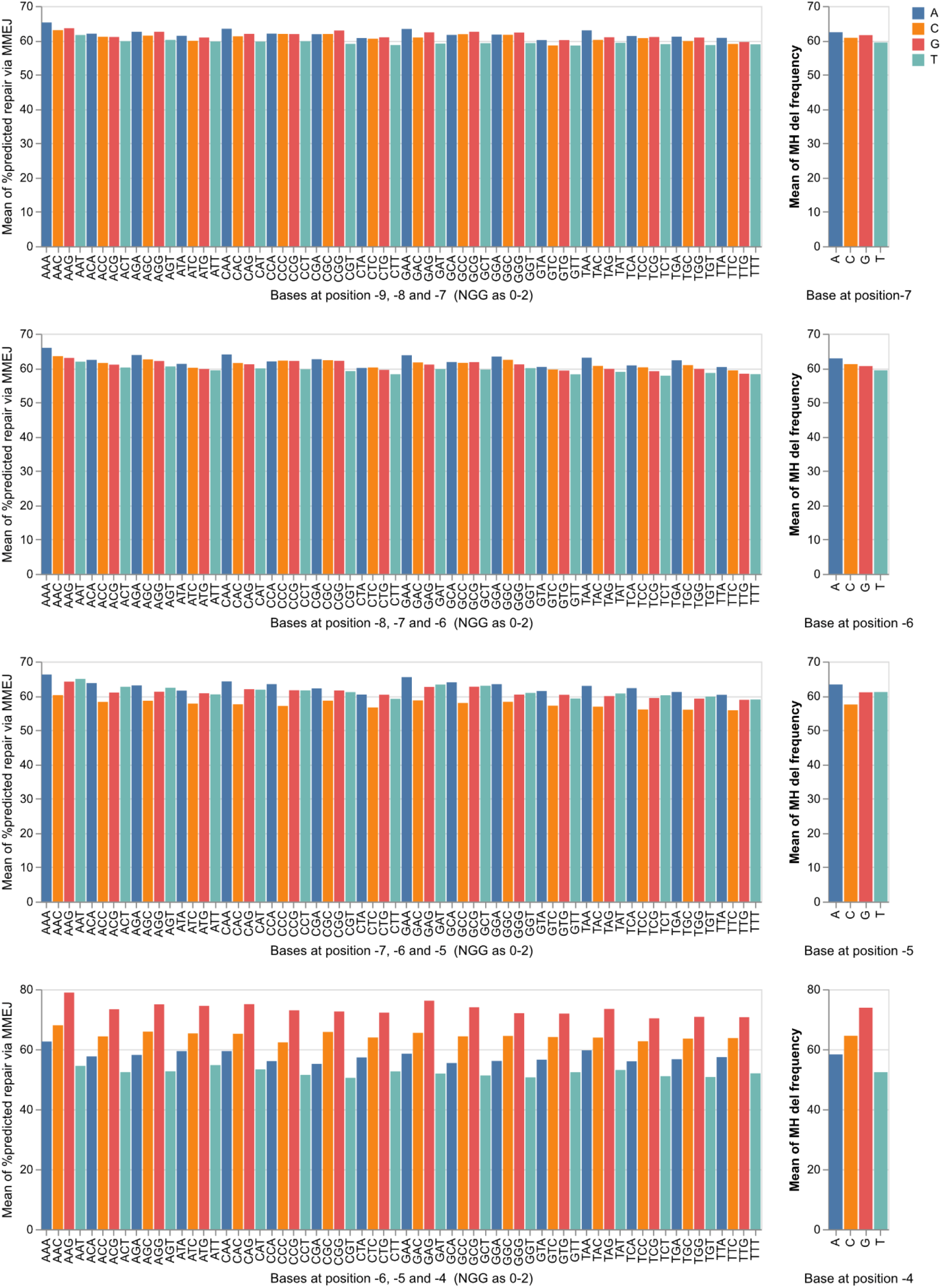
Modelling the predicted percentage of repair via MMEJ based on the base-composition of the gRNA binding site in the genome reveals that base composition at position -4 increases the % of gene editing outcomes by MMEJ for gRNAs with PAM “NAA”. We performed an analysis across a subset of the human coding exome using the InDelphi-HEK293 model retrieving 1,751,128 unique gRNAs. A subset of 500,00 gRNAs was randomly selected for plotting. This reveals that there is an enrichment for expected % of gene editing outcomes by MMEJ when the base at position -4 is a C or G, when compared to A or T. Such enrichment is not apparent for the positions -5 through -9 (counting the NGG PAM as nucleotides 0-2).

**Supplementary figure 6:**
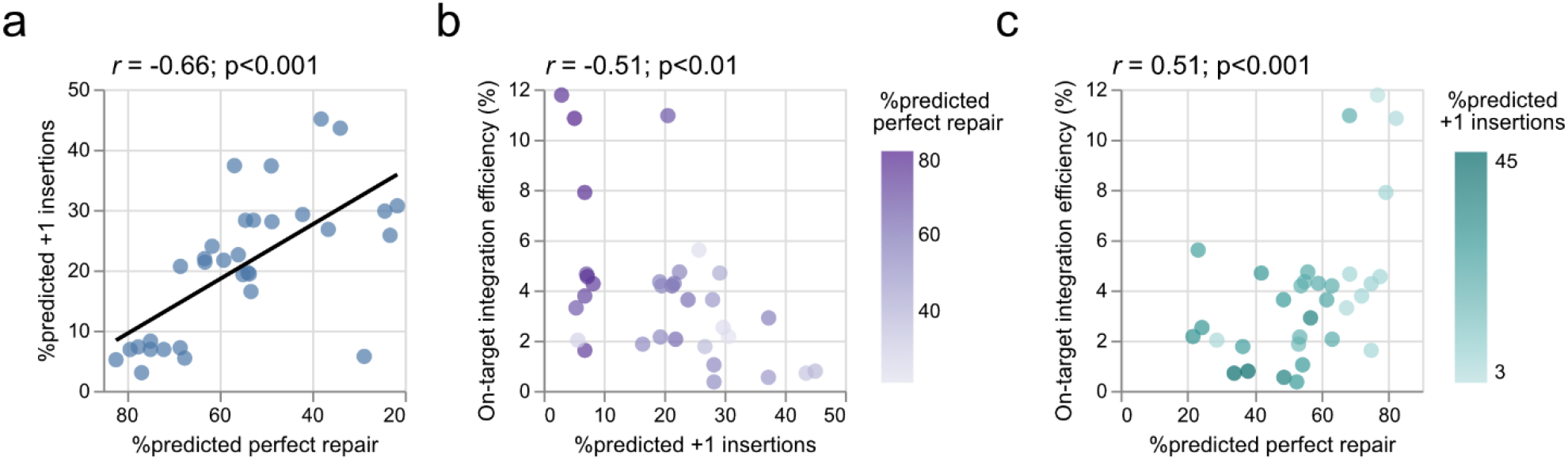
Correlating experimental on-target integration efficiencies to InDelphi predictions. **(a)** There is an inverse correlation between the % of predicted perfect repair (defined as any repair mobilizing a trimology repeat) and the % of predicted +1 insertions. (b-c) As such, on-target integration efficiencies decrease when there is a higher % of prediction for +1 insertions and increase when there is a higher % of predicted perfect repair.

**Supplementary figure 7:**
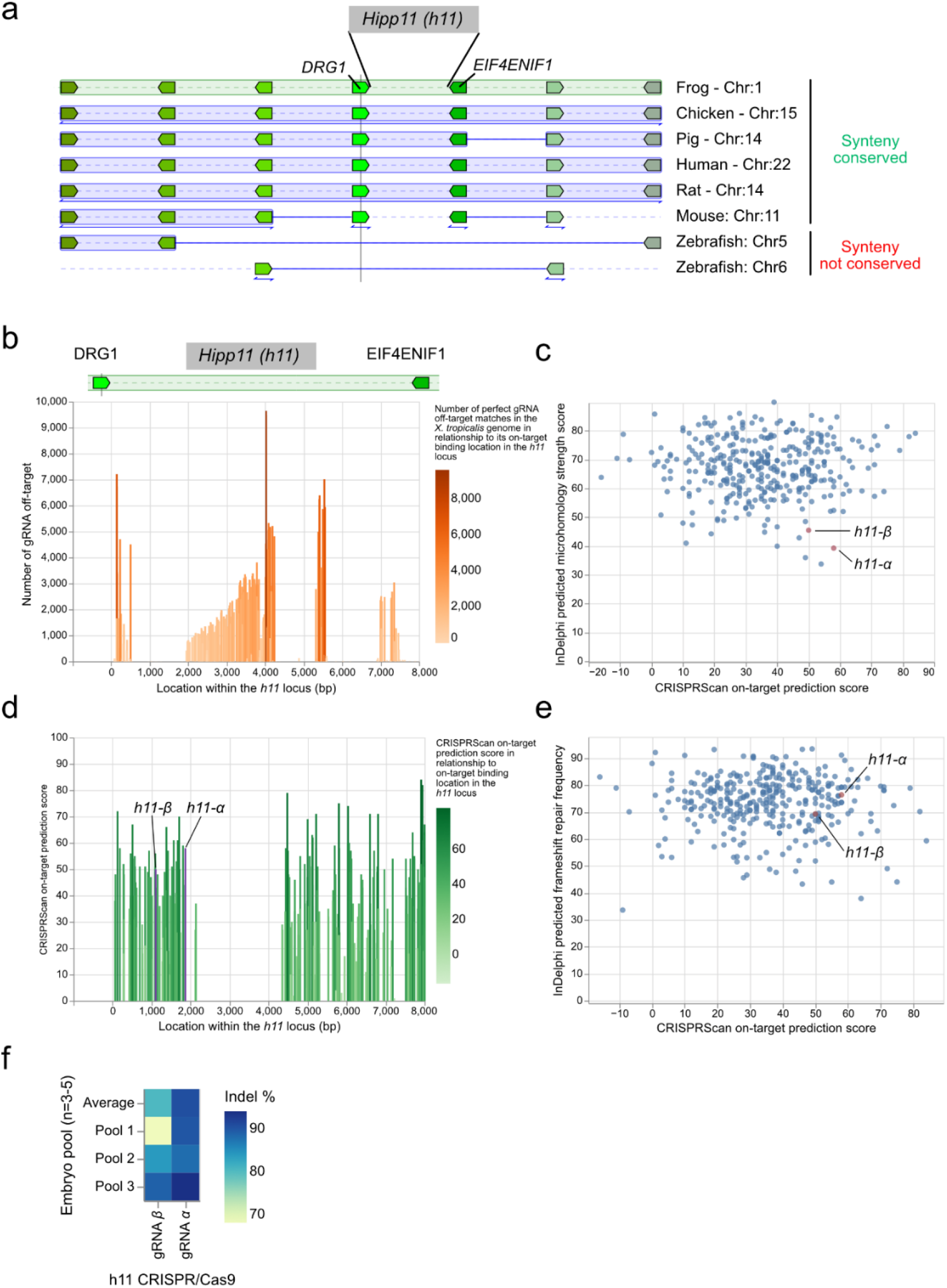
The *Xenopus tropicalis hipp11* locus can be gene edited by CRISPR/Cas9. **(a)** Using genome synteny (Genomicus^35^), we identify conservation of the intergenic region between *drg1* and *eif4enif1* corresponding to the *Hipp11* (*H11*) transgene landing site previously described in human, mouse and pig. Of note, synteny here is not conserved in the teleost lineage. **(b)** 55.3% (447 gRNAs/809) of potential gRNAs targeting this region have an unacceptable off-target profile with >1 perfect off-target match in the *Xenopus tropicalis* genome. **(c-e)** For the remaining gRNAs, we calculated via InDelphi-mESC the predicted frequency of editing outcomes via MMEJ, predicted frameshift repair frequency and the on-target CRISPRScan score to identify two suitable gRNA target sites (*h11-α* and *h11-β*) spaced 767bp apart. **(f)** One-cell stage *X. tropicalis* embryos were injected with CRISPR/Cas9 targeting either *h11-α* or *h11-β*. Embryos were grown until stage 23, lysed (pools of 3 to 5 embryos) and genome editing efficiencies are shown, as determined by sanger sequencing and trace deconvolution.

**Supplementary figure 8:**
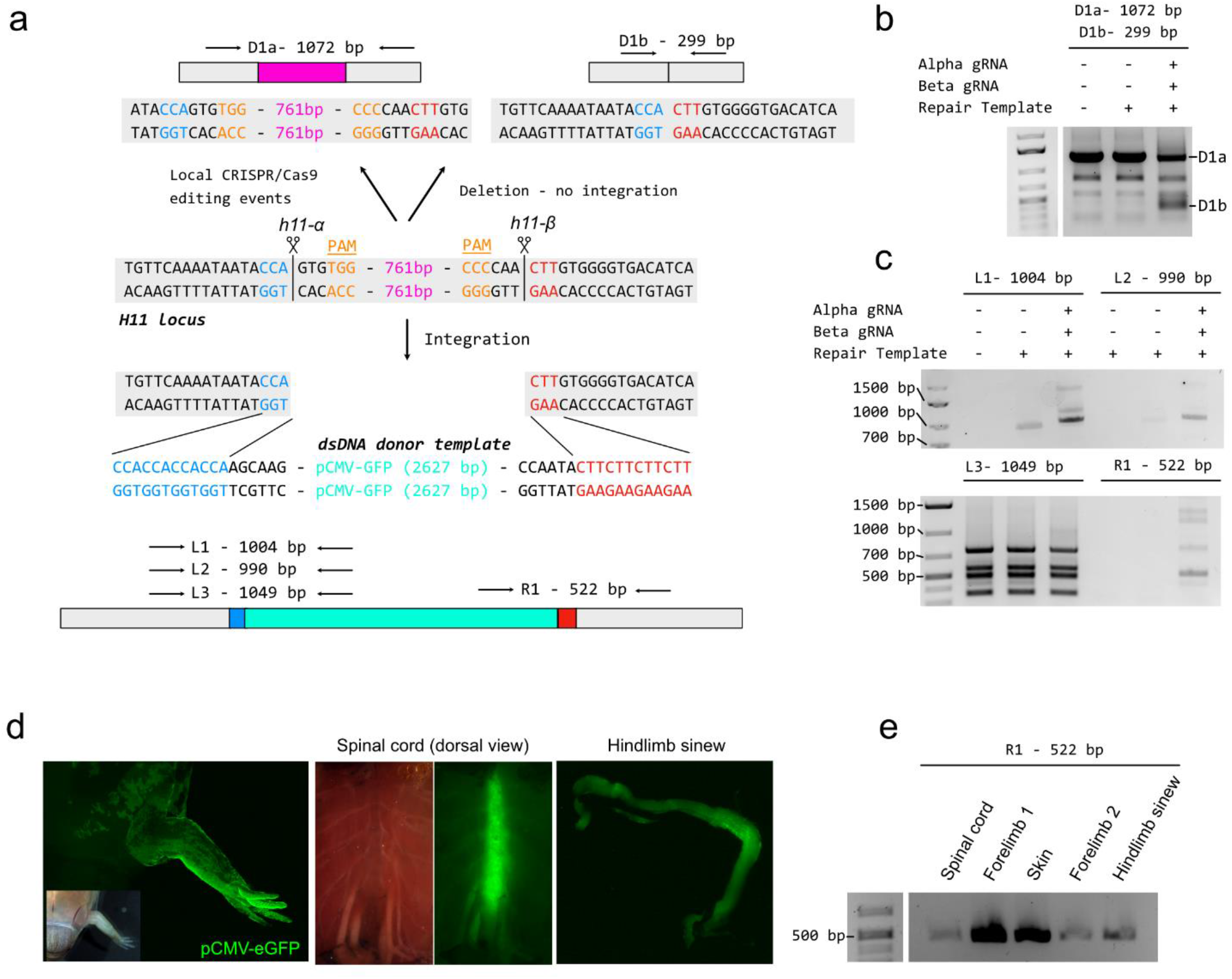
Stable integration into the *X. tropicalis hipp11* locus. **(a)** Schematic of the CRISPR/Cas9 integration strategy. Co-delivering two RNPs (*h11-α* and *h11-β*) together with dsDNA donor template leads to several potential gene editing outcomes. Firstly D1a, amplified are unedited and locally edited (Indels) genome copies. Secondly D1b, amplified in case of CRISPR-mediated deletion of intervening DNA between the binding site of *h11-α* and *h11-β*. Thirdly, integration of the dsDNA donor template, in place of the intervening DNA between the binding site of *h11-α* and *h11-β*. **(b)** Co-delivering *h11-α* and *h11-β* RNP results in a lower amplification of product D1a, and an increase in product D1b, showcasing CRISPR-mediated deletion as an editing outcome. All lanes are pools of 50 injected embryos. **(c)** 5’ (L1, L2, L3) and 3’ (R1) junction products can be amplified, revealing targeted integration of pCMV-eGFP in the *hipp11* locus. **(d)** Example of clonal expansion of GFP+ cells as an embryo develops from neurula to metamorphosis. **(e)** Several GFP+ tissues were dissected, lysed and the 3’ (R1) junction products amplified, revealing stable long-term integration into the *hipp11* locus

**Supplementary figure 9:**
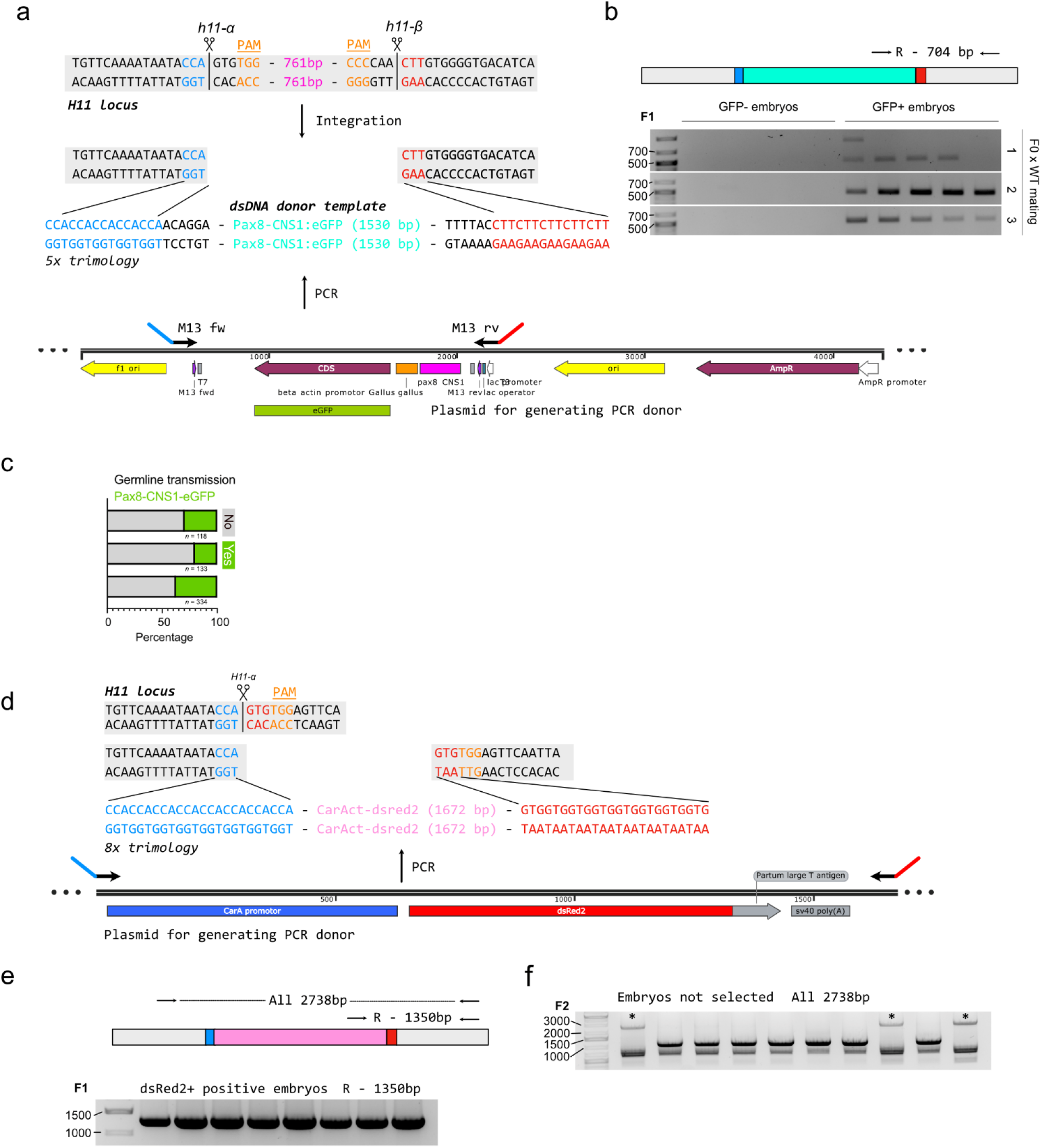
Stable integration into and tissue-specific expression from the *X. tropicalis hipp11* locus. **(a)** Schematic CRISPR/Cas9 integration strategy. Co-delivering two RNPs (*h11-α* and *h11-β*) together with Pax8-CNS1:eGFP dsDNA donor template (5x trimology) generated by simple overhang PCRs from a plasmid **(b)** In F1 offspring, 3’ (R) junction products were amplified, revealing targeted integration of Pax8-CNS1:eGFP in the *hipp11* locus and positive germline transmission. **(c)** In six F0 founder animals, three (50%) demonstrated transmission through germline at mean rate of 29.1% ± 8.6%. **(d)** Schematic CRISPR/Cas9 integration strategy. Delivering two RNPs (*h11-α*) together with CarAct:dsRed2 dsDNA donor template (8x trimology) generated by simple overhang PCRs from a plasmid. **(e)** In F1 offspring, 3’ (R) junction products were amplified, revealing targeted integration of Pax8-CNS1:eGFP in the *hipp11* locus and positive germline transmission. **(f)** F2 homozygote knock-in animals (stars) can be identified by a PCR with primers binding the genomic sites left and right of *h11-α*, thus amplifying the entire fragment (2738 bp) in homozygotes knock-in animals.

**Supplementary Figure 10:**
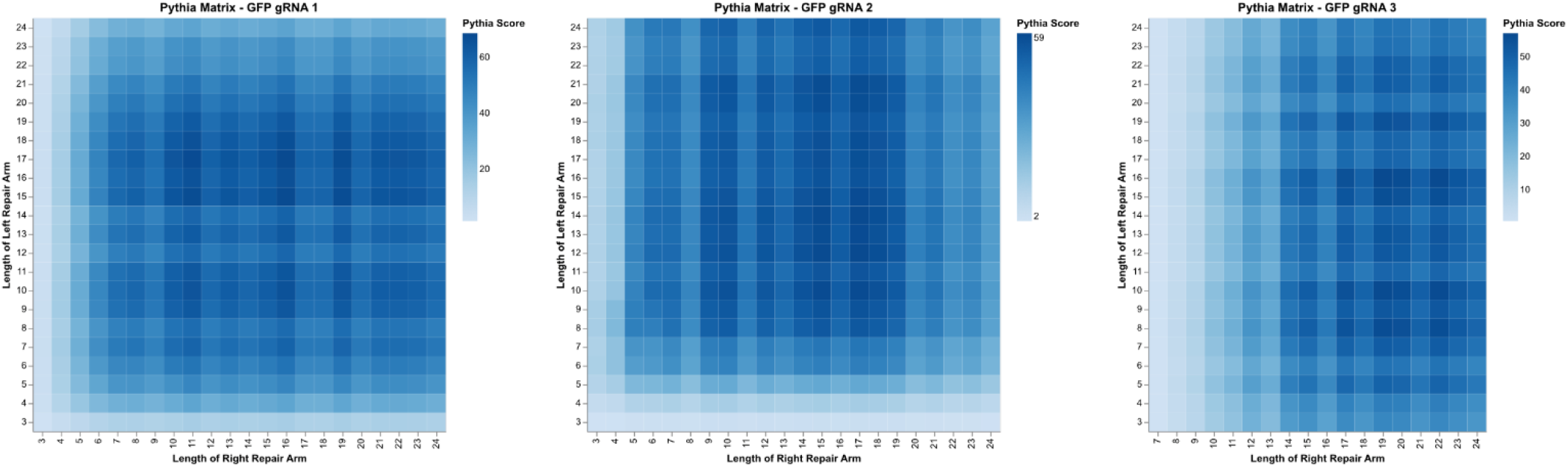
Pythia Matrices for GFP gRNAs.

**Supplementary Figure 11:**
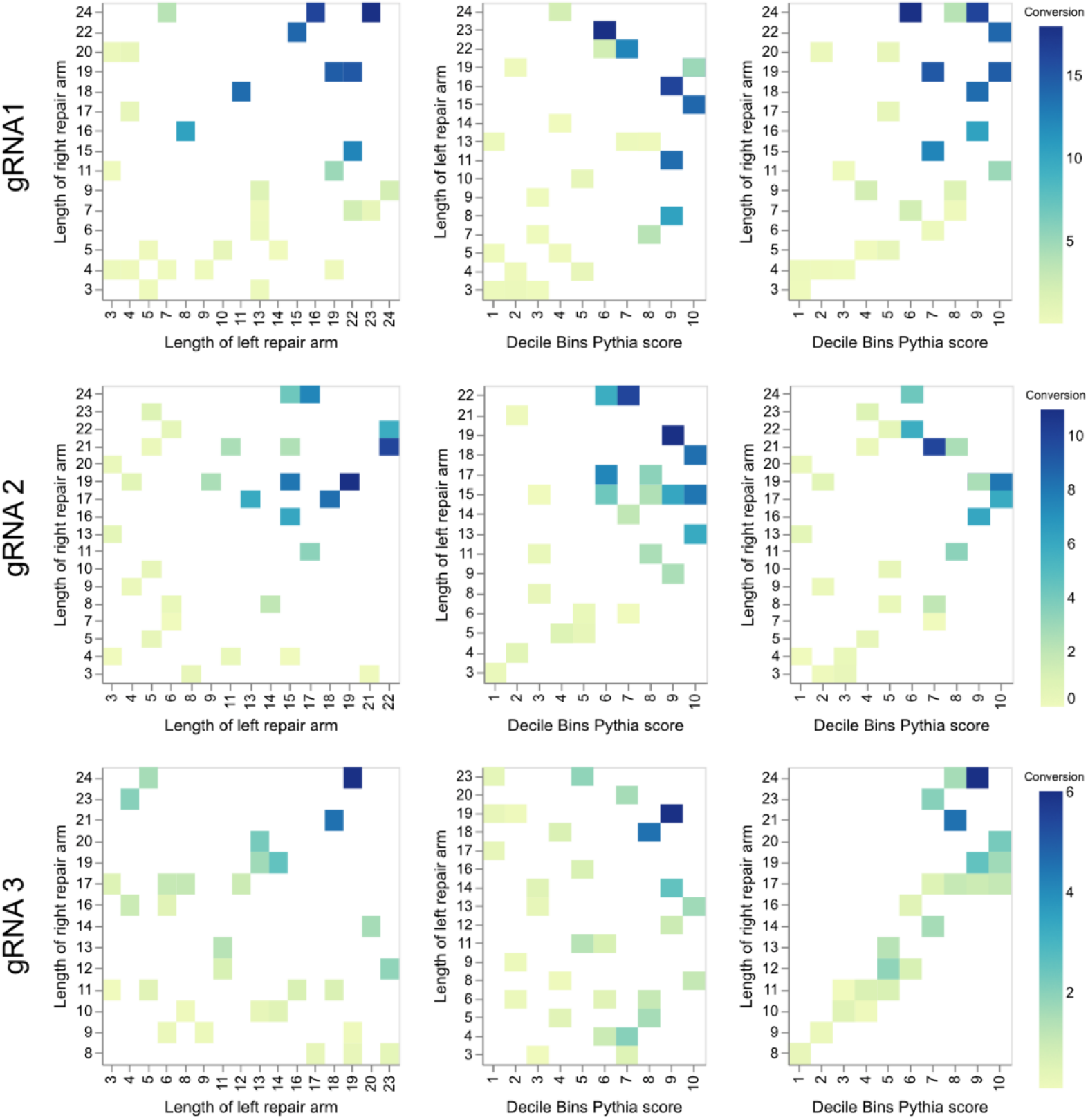
Relationship between lengths of repair arms, Pythia scores and eGFP-to eBFP conversion efficiencies. Each row represents one GFP gRNA (top 1, middle 2, bottom 3) each with 30 distinct repair templates binned across Pythia prediction score (predicted %perfect repair). Both left and right repair arm length is related to conversion (eGFP-to eBFP) efficiencies. Further, there is a direct correlation between Pythia prediction scores and conversion efficiencies.

### Supplementary Movies

**Supplementary Movie 1:** Benchtop mesoSPIM recording of an adult kidney from an F0 *Xenopus tropicalis* with a stable integration of pax8-CNS1:eGFP in the *hipp11* stable landing site.

**Supplementary Movie 2:** Time-lapse imaging of tadpole development in a stable F1 *Xenopus tropicali*s line with a stable integration of pax8-CNS1:eGFP in the *hipp11* stable landing site.

**Supplementary Movie 3:** Time-lapse imaging of tadpole development in a stable F1 *Xenopus tropicali*s with a stable integration of CarAct:dsRed2 in the *hipp11* stable landing site.

**Supplementary Movie 4:** Benchtop mesoSPIM and Schmidt objective two-photon microscopy recording of a stable F2 *Xenopus tropicali*s with a stable integration of CarAct:dsRed2 in the *hipp11* stable landing site.

**Supplementary Movie 5:** Benchtop mesoSPIM recording of wildDisco processed adult mouse brain demonstrating viral delivery and in-frame eGFP-tagging of *Tubb2a*.

### Supplementary Tables

**Supplementary Table 1:**
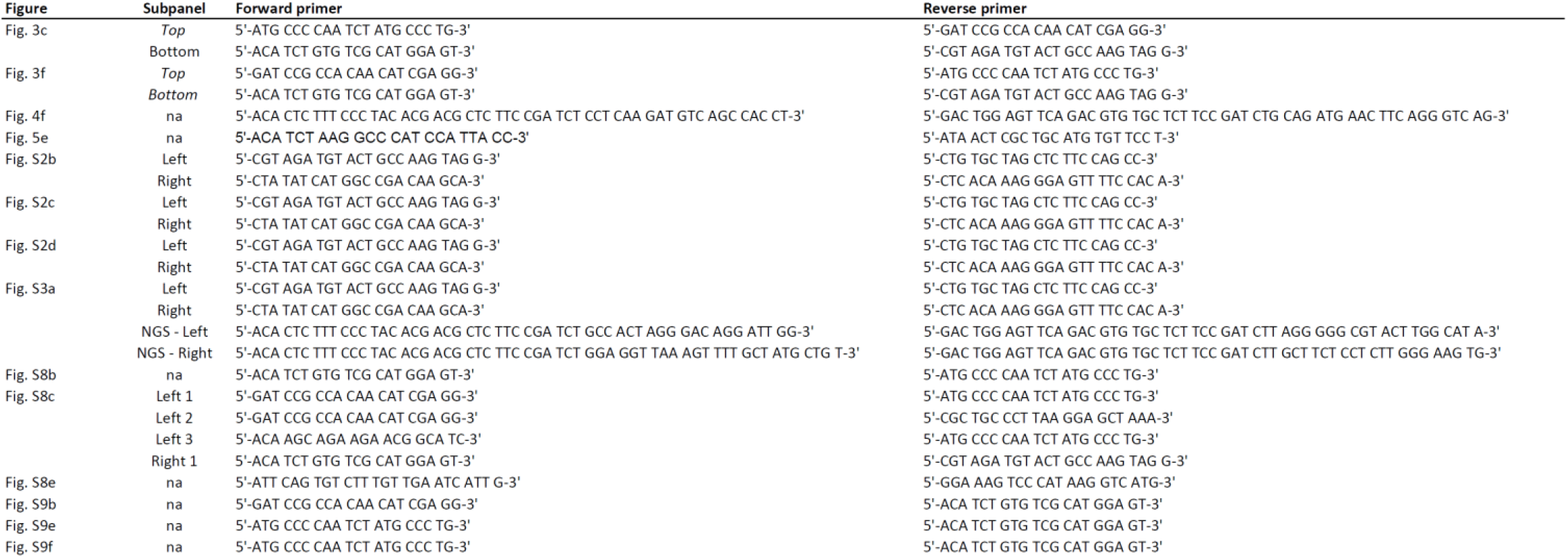
Primer sequences used for PCR amplification.

**Supplementary Table 2:**
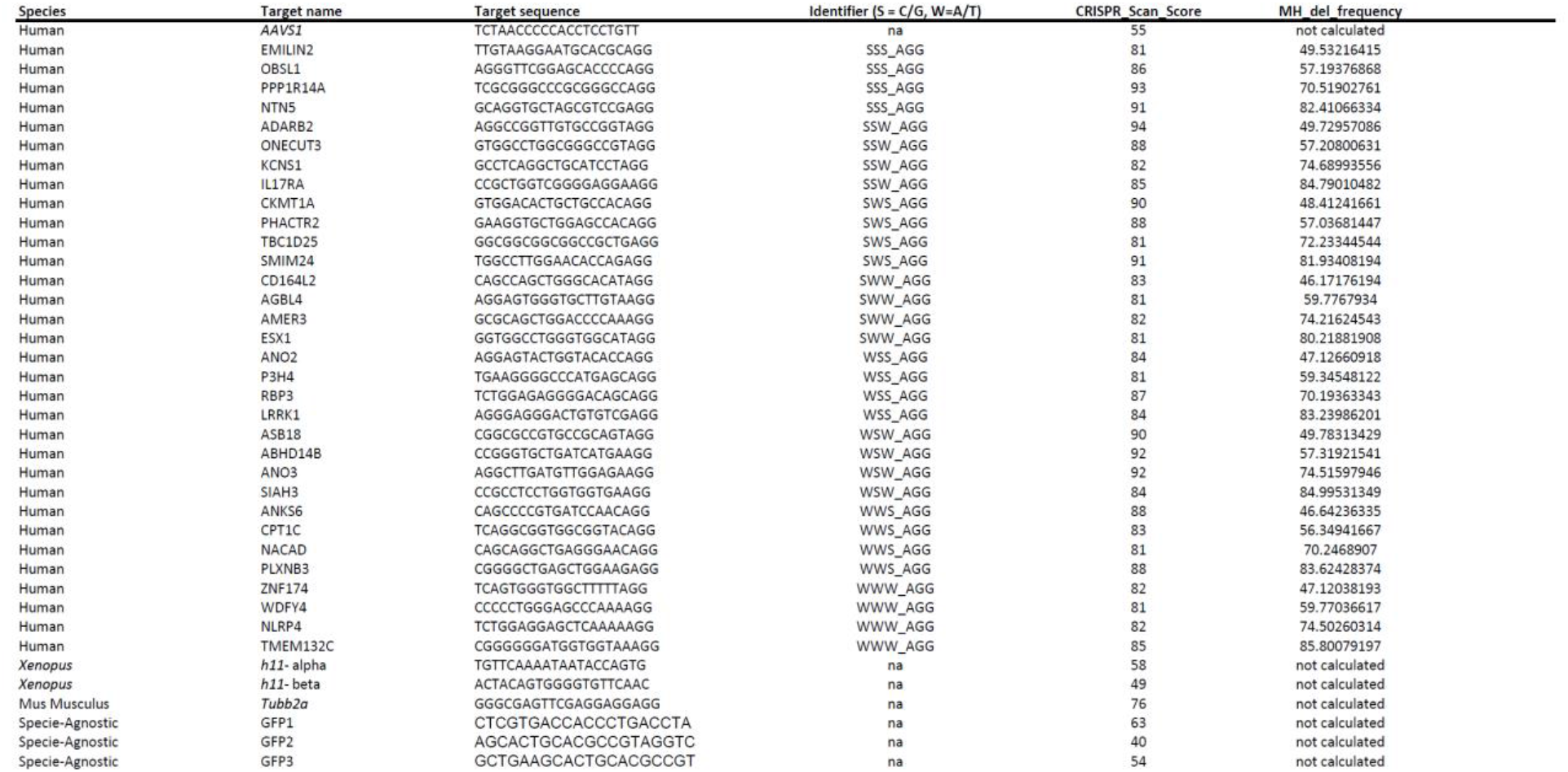
Sequences, CRISPRScan scores and InDelphi metrics for gRNAs.

**Supplementary Table 3:**
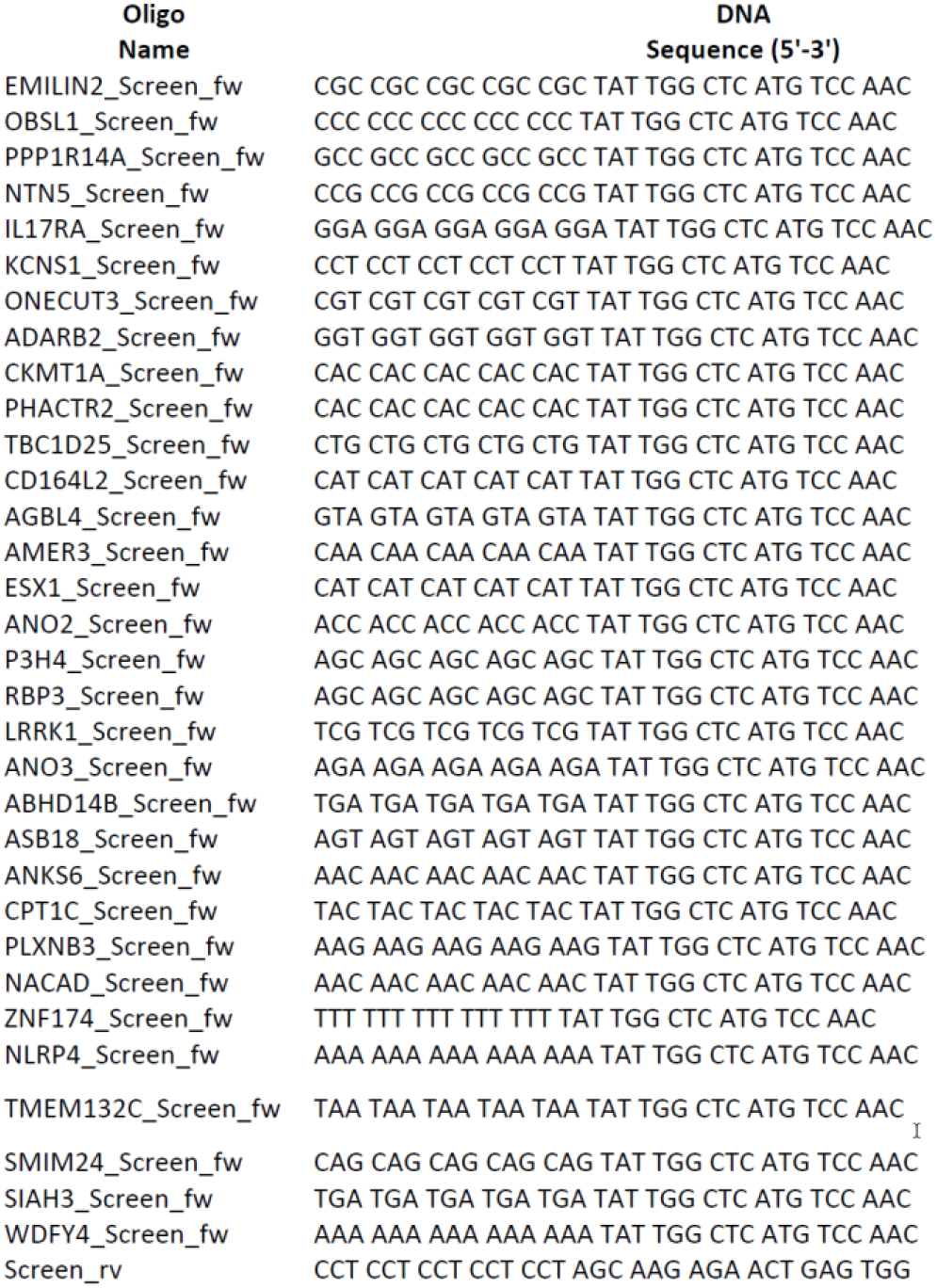
Primer sequences used for amplification of pCMV-eGFP repair cassette from AAV-CMV-GFP (addgene #67634) for each of the 32 target loci.

**Supplementary Table 4:**
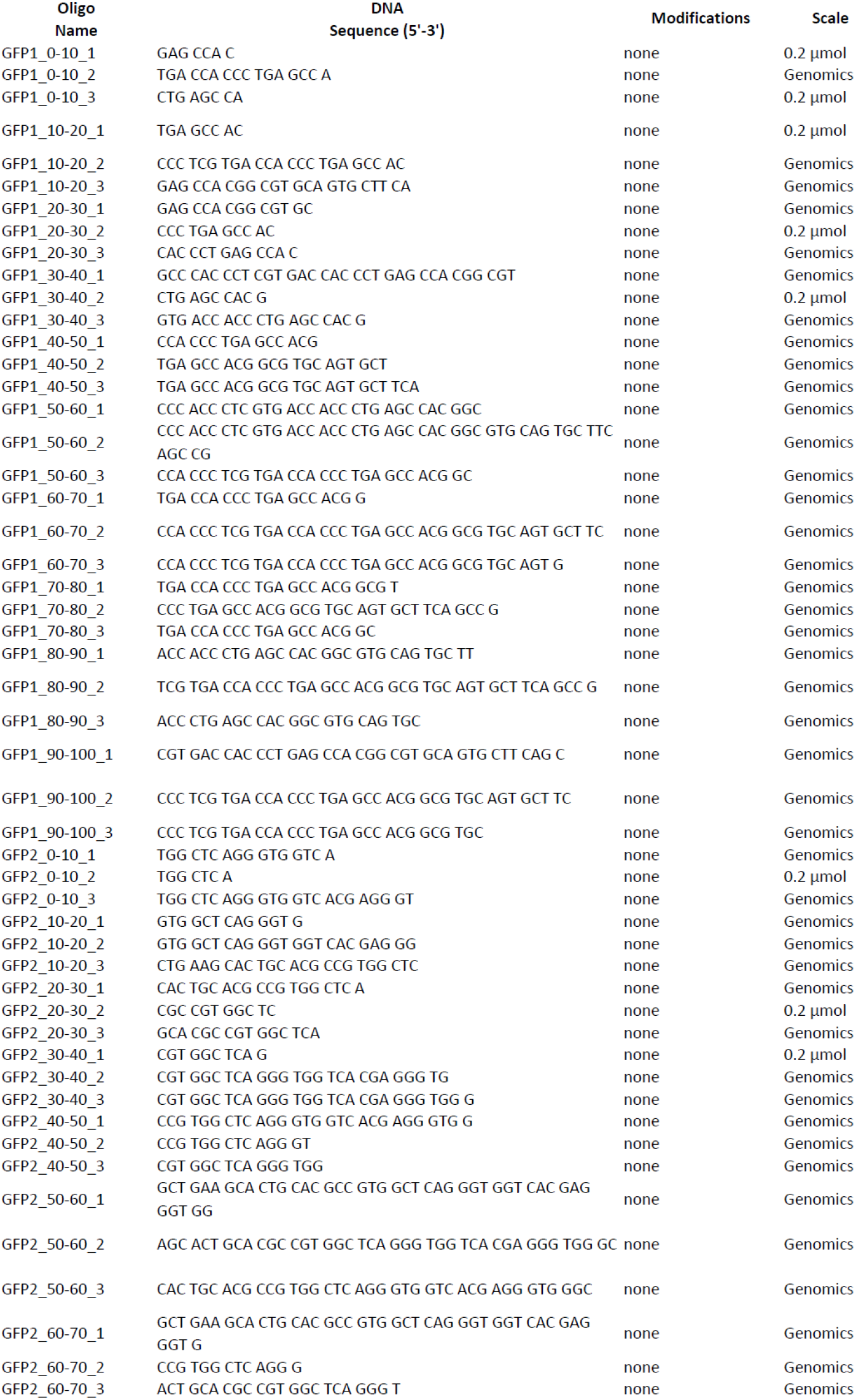
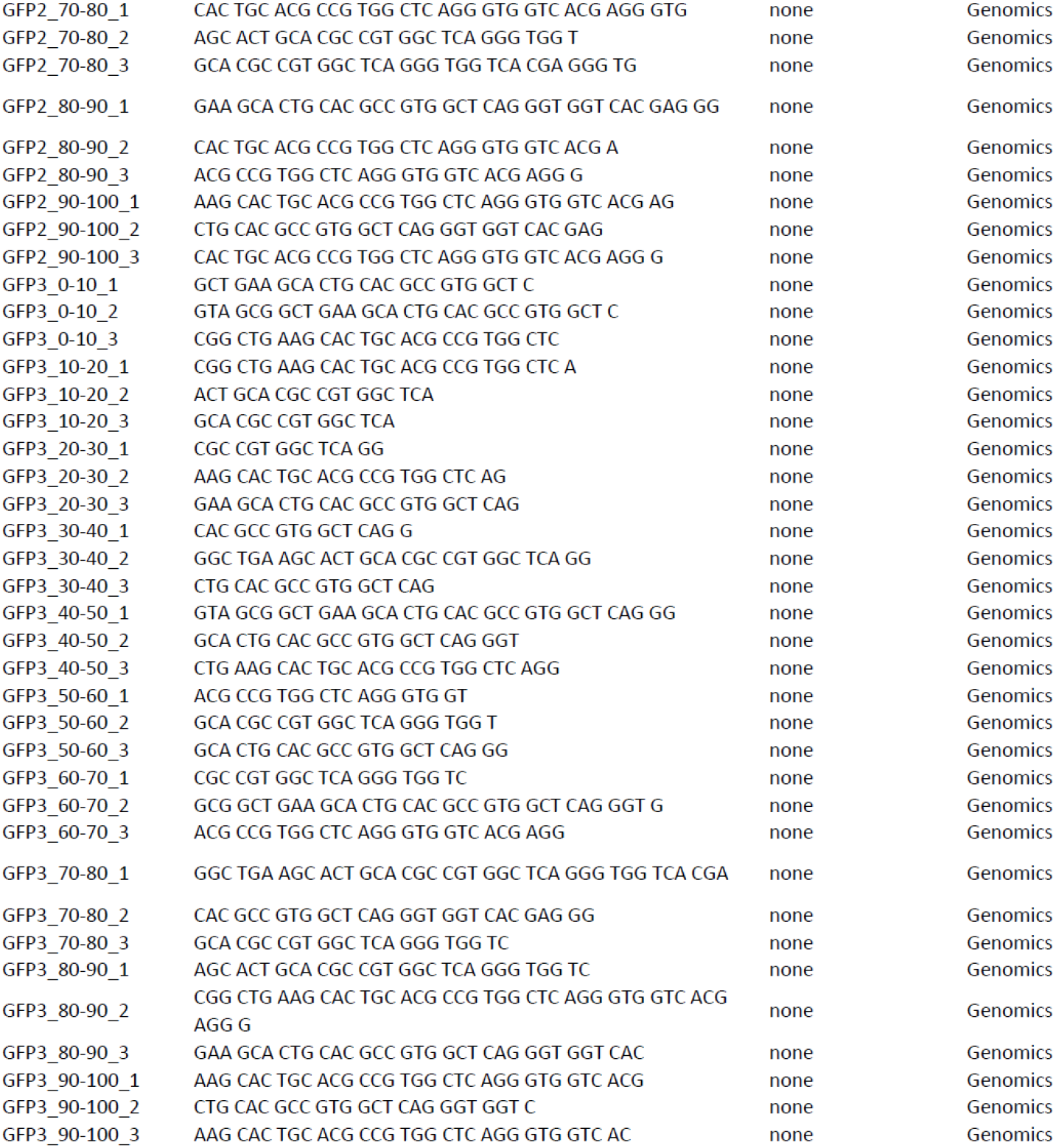
Sequences of ssODN repair templates used for eGFP -> eBFP conversion in HEK293T cells.

## Notes

http://www.pythia-editing.org/

